# Carotenoid pigments enhance rhodopsin-mediated phototrophy by light-harvesting and photocycle-accelerating

**DOI:** 10.1101/2024.11.08.622755

**Authors:** Takayoshi Fujiwara, Toshiaki Hosaka, Masumi Hasegawa-Takano, Yosuke Nishimura, Kento Tominaga, Kaho Mori, Satoshi Nishino, Yuno Takahashi, Tomomi Uchikubo-Kamo, Kazuharu Hanada, Takashi Maoka, Shinichi Takaichi, Keiichi Inoue, Mikako Shirouzu, Susumu Yoshizawa

**Author notes:** Correspondence (K.I.), (M.S.), (S.Y.). These authors contributed equally: Takayoshi Fujiwara and Toshiaki Hosaka. Senior authors (Keiichi Inoue, Mikako Shirouzu, Susumu Yoshizawa).

## Abstract

Microbial rhodopsins, photoreceptor proteins widely distributed in marine microorganisms, receive large amount of light energy that sustains marine ecosystems. Although rhodopsins generally harbor retinal as their only chromophore, a recent study reported that carotenoid antennae transfer light energy to the retinal in proteorhodopsins, proton pump rhodopsins abundant in marine environments. Here, using marine bacterial isolates, we detected energy transfer from a carotenoid (myxol) to retinal not only in proteorhodopsin but also in the chloride ion-pumping rhodopsin. Carotenoid binding improved the light utilization efficiency of the proteorhodopsin by accelerating the photocycle, together with facilitating light-harvesting. The carotenoid-binding ability is conserved in rhodopsins of the phylum *Bacteroidota*, which are widely transcribed in the photic zone. These findings suggest that the distribution of carotenoid-binding rhodopsins is more taxon-specific than previously thought, thus underscoring the importance of carotenoid-binding rhodopsins that provide an extended light utilization strategy in the environmental adaptation of marine bacteria.

## Introduction

Marine ecosystems are maintained depending on the energy provided by sunlight. It was thought that light energy flows into marine ecosystems in the form of organic matter, mostly through the carbon fixation process of photosynthesis. However, the discovery of proteorhodopsin (PR), a light-driven H^+^ pump abundant in marine bacteria, has revealed an alternative pathway for light energy to flows into the ecosystems^1^. Microbial rhodopsins (hereafter referred to as rhodopsins) are widely distributed photoreceptor proteins in bacteria, archaea, and eukaryotes^2,3^, using retinal as their chromophore. In addition to PR, several other rhodopsins have been identified in marine bacteria with different ion transport activities, including sodium-ion and chloride-ion pumps^4,5^. It has been estimated that over half of the marine bacteria inhabiting the ocean surface possess rhodopsin genes^6^, and that these bacteria receive light energy comparable to the amount absorbed by photosynthesis^7^.

Xanthorhodopsin (XR), a light-driven proton pump, discovered in the extremely halophilic bacterium *Salinibacter ruber* M31^T^, exceptionally utilizes two chromophores: an all-*trans* retinal and the 4-keto carotenoid pigment (salinixanthin), for enhancement of light-harvesting efficiency^8,9^. The salinixanthin is located on the fenestration which exposes the retinal-binding pocket in the protein to the outside, with the ionone-rings of the two chromophores positioned just 5 Å apart^10^. This unique arrangement allows for efficient excitation energy transfer (EET) between the chromophores, the salinixanthin functions as an antenna, and thus XR receives a broader range of wavelengths (560 nm from the retinal chromophore and 450–520 nm from the salinixanthin)^8,11^. A recent study discovered that hydroxylated carotenoid pigments (zeaxanthin and lutein) also work as antennae for PRs in marine bacteria, namely KR1 (found in *Dokdonia eikasutus* NBRC 100814^T^) and TsPR (found in *Tenacibaculum* sp. SG-28)^12^, revealing that carotenoid antennae are not limited to rhodopsins in extreme environments, but also exist in marine environments.

The marine-dominant phylum *Bacteroidota*, which includes the genera such as *Dokdonia* and *Tenacibaculum*, is known to produce a wide variety of carotenoids. This suggests that the interactions between their rhodopsins and intracellular carotenoids may not be uniform^13^. However, the roles of carotenoids beyond their functioning as light-harvesting antennae is not known, and attempts to find antennae-bound rhodopsins that transport ions other than H^+^ have been unsuccessful so far^14^. In this study, we used *Nonlabens marinus* S1-08^T^, a member of the marine *Bacteroidota* that carries three different types of rhodopsin genes^4^, to explore the mechanisms by which carotenoids enhance rhodopsin function. By co-incubating rhodopsins and carotenoids extracted from S1-08^T^ cells, we demonstrated the binding ability of these rhodopsins to carotenoids. Spectroscopic and structural analyses of rhodopsin–carotenoid complexes were conducted to examine the effects of carotenoid binding to rhodopsins. Moreover, phylogenetic and metatranscriptomic analyses revealed that the carotenoid-binding PRs are conserved among marine *Bacteroidota* and widely transcribed under various environmental conditions in the photic zone.

## Results

### Three rhodopsins in *N. marinus* S1-08^T^ bind to intracellular carotenoids

Energy transfer from carotenoids to the retinal chromophore requires a fenestration structure in rhodopsin, which is facilitated by a glycine residue in transmembrane domain 5 (TM5; Gly-156 in XR)^15^. The small side chain of glycine residue at this position provides sufficient space for the 4-keto ring of salinixanthin in XR, and the hydroxyl ring of zeaxanthin in Kin4B8 (Xanthorhodopsin-like rhodopsin, XLR clade)^12,15^. In contrast, in bacteriorhodopsin from *Halobacterium salinarum* (HsBR), where the glycine residue is replaced by the bulky residues, Trp-138, the fenestration structure cannot form, and thus it cannot use a carotenoid antenna^15^. To explore rhodopsins with the ability to bind carotenoids, we aligned and compared amino acid sequences of rhodopsins (Fig. S1). A phylogenetic tree and multiple sequence alignment revealed that this glycine residue was conserved in many rhodopsins belonging to PR, XLR, chloride ion-pumping rhodopsin (ClR), and sodium ion-pumping rhodopsin (NaR) clades (Fig. 1A, 1B, and Fig. S1).

**Figure 1.**
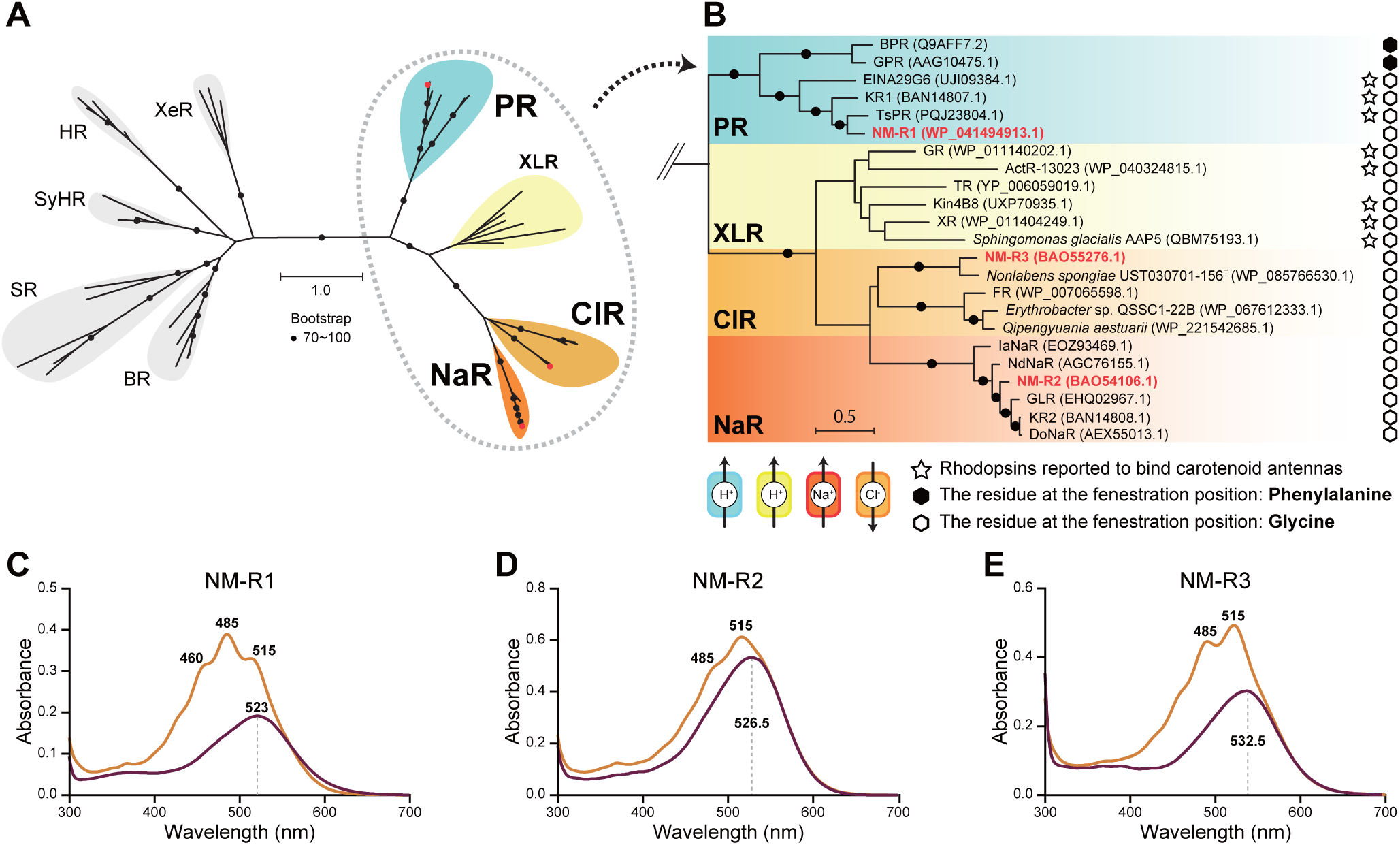
Three types of rhodopsins from *Nonlabens marinus* S1-08^T^ bind to carotenoids extracted from S1-08^T^ cells. (A) Phylogenetic distribution of antenna-containing rhodopsins. Bootstrap probabilities (≥ 70%) are indicated by circles. (B) Enlarged view of rhodopsins from *N. marinus* S1-08^T^. (C–E) Absorbance spectra of purified NM-R1, NM-R2, and NM-R3 before (purple) and after (orange) adding whole carotenoids extracted from S1-08^T^ cells (NmC), respectively.

Following this result, we conducted an experiment to detect the rhodopsin– carotenoid complex in the marine bacterium *N. marinus* S1-08^T^, a member of *Bacteroidota*^12^. *N. marinus* S1-08^T^ carries three distinct rhodopsin genes: NM-R1 (*N. marinus* rhodopsin 1; PR clade), NM-R2 (*N. marinus* rhodopsin 2; NaR clade), and NM-R3 (*N. marinus* rhodopsin 3; ClR clade), and produces carotenoids^4^. Purified His-Tag fusion rhodopsin proteins, heterologously expressed in *E. coli* cells, were mixed with carotenoids extracted from S1-08^T^ cells (hereafter NmC) and purified again. After the second purification, two bands at 485 and 515 nm appeared in the absorption spectra of all rhodopsins (Fig. 1C-E), confirming that NM-R1, NM-R2, and NM-R3 were capable of binding carotenoids produced by S1-08^T^. Further spectroscopic analyses focused on NM-R1 and NM-R3, which exhibited the higher level of carotenoid binding.

To identify the carotenoids bound to rhodopsins, the molecular structures of carotenoids produced by S1-08^T^ were analyzed. High-performance liquid chromatography with diode array detector (HPLC-DAD) elution profile of NmC exhibited two major peaks at 7.1 and 8.6 min (Fig. 2A). These carotenoids were identified as (3*R*,2’*S*)-myxol and (3*R*,3’*R*)-zeaxanthin, respectively, based on the spectroscopic data of absorption spectra by DAD, mass spectra by LC-MS, ^1^H-NMR spectra, and circular dichroism (CD) spectra (Fig. 2B, 2C, and Table S1). The HPLC-DAD elution profile of the carotenoids extracted from the rhodopsin–carotenoid complexes showed that the chromophores of NM-R1 consisted of both zeaxanthin and myxol, whereas the chromophores of NM-R3 mainly consisted of myxol (Fig. 2A and 2C). These pigment analyses demonstrated that NM-R1 bound to both myxol and zeaxanthin, NM-R3 selectively bound to myxol.

**Figure 2.**
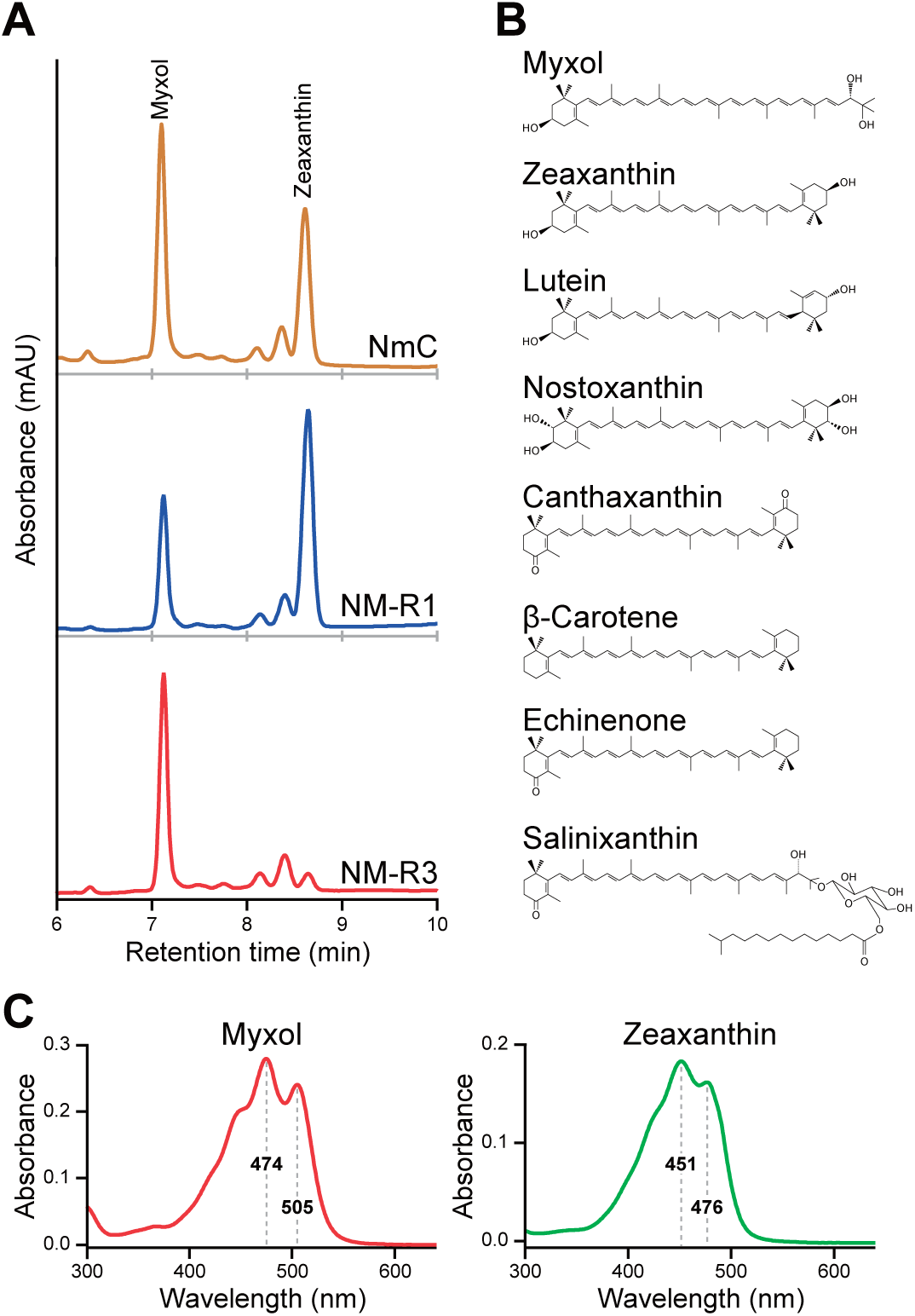
Identification of carotenoid pigments. (A) HPLC profile of carotenoid pigments, registered at 450 nm. The red, orange, and blue chromatograms are NmC, binding carotenoid pigments to NM-R1, and binding carotenoid pigments to NM-R3, respectively. (B) 2D structures of myxol and known carotenoid pigments that act as antennae. (C) Absorbance spectra of myxol and zeaxanthin in MOPs buffer.

### The bound carotenoids in NM-R1 and NM-R3 interact with retinal

To confirm the, interaction between the carotenoids and retinal, one approach is to measure the differential absorbance spectrum before and after bleaching with hydroxylamine (HA)^8^. HA hydrolyses the Schiff base bond which connects the retinal chromophore to the rhodopsin protein^16^. When the NM-R1 and NmC mixture was incubated with HA, the resulting the difference spectrum showed a broad peak at 540 nm, reflecting the hydrolysis of the retinal Schiff base, and peaks at 363 nm due to retinal oxime formation, along with peaks at 452, 485, and 520 nm, similar to XR (Fig. 3A)^17^. These peaks were thought to originate from carotenoid molecules interacting with the retinal, altering their absorbance spectra upon the formation of the complex^8^. Another method is measuring the CD spectrum of the rhodopsin–carotenoid complex. For example, the CD spectrum of XR–salinixanthin complex exhibits a negative band from the retinal chromophore and positive bands from the bound salinixanthin^18^. However, in the CD spectrum of archaerhodopsin-2–bacterioruberin, which lacks antenna function, positive bands from both chromophores are observed^18,19^. The CD spectrum of NM-R1– NmC complex exhibited several bands in the visible wavelength region, at 453 (+), 477 (+), and 531 nm (−), supporting the specific interaction between NM-R1 to NmC (Fig. 3B). Furthermore, the same experimental analyses were conducted on mixtures or complexes of NM-R1 to carotenoids extracted from *Tenacibaculum* sp. SG-28 cells (producing only zeaxanthin, hereafter TsC), and *N. spongiae* JCM 13191^T^ cells (producing only myxol, hereafter NsC)^12,20^. The differential spectra upon HA hydrolysis reaction and the CD spectra of both samples were different with each other, indicating that NM-R1 can interact with bound zeaxanthin and myxol (Fig. 3C-F). Caused by HA bleaching, the difference spectrum of NM-R3 and NmC mixture exhibited bands at 470, 500, and 530 nm (Fig. 3G). The CD spectrum of NM-R3–NmC complex also exhibited positive and negative bands in the visible wavelength region at 461 (+), 485 (+), and 541 nm (−), suggesting a specific interaction between NM-R3 to NmC (Fig. 3H).

**Figure 3.**
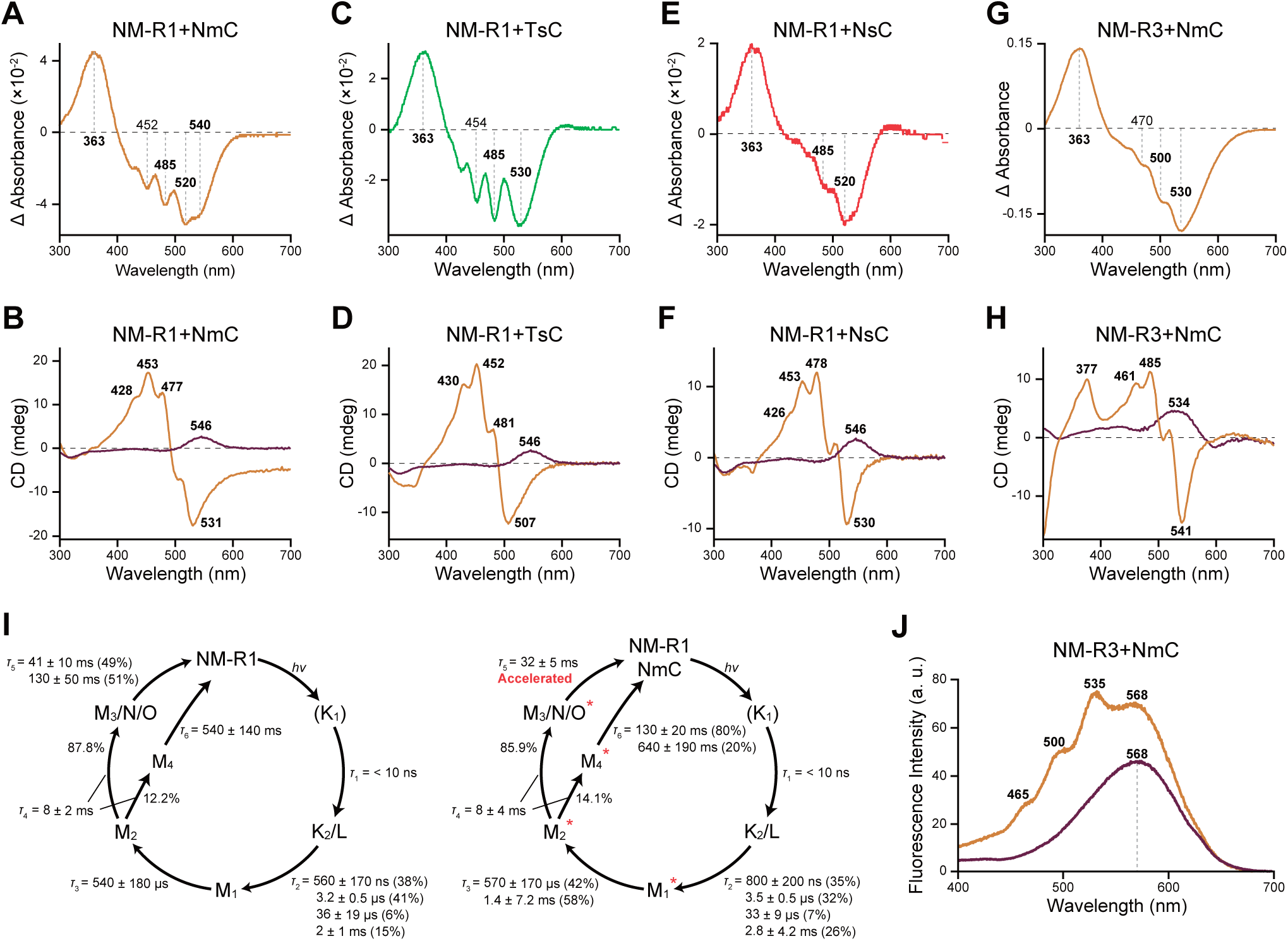
Spectroscopic analyses of NM-R1 and NM-R3. (A, C, E) Absorbance changes of NM-R1 with NmC (myxol and zeaxanthin), TsC (zeaxanthin), and NsC (myxol), caused by incubation with hydroxylamine for 1 h, 30 min, and 30 min, respectively. (B, D, F) Circular dichroism spectra of NM-R1 with (orange) or without (purple) NmC, TsC, and NsC, respectively. (G) Absorbance changes of NM-R3 with NmC, caused by incubation with hydroxylamine for 9 h. (H) Circular dichroism spectra of NM-R3 with (orange) or without (purple) NmC. (I) Photocycle model of NM-R1 without (left) and with (right) NmC. The percentages of the pre-exponential factors for each component with the indicated lifetimes of the multi-exponential processes. The photointermediates whose absorption spectra included the absorption changes of carotenoid are indicated by red asterisks. (J) Fluorescence excitation spectra of NM-R3 with (orange) and without (purple) myxol; emission was monitored at 720 nm.

### Carotenoid binding enhances rhodopsin functions in the two ways: as light-harvesting antennae and as photocycle-accelerating pigments

When the retinal chromophore receives light energy and undergoes isomerization from the all-*trans* to the 13-*cis* form, the rhodopsin protein produces several photointermediates and finally returns to its original state. This series of photochemical reactions is referred to as the photocycle. To investigate how bound carotenoids affect this process, transient absorption spectra of rhodopsins were monitored. The transient absorption measurements of NM-R1 dominantly detected the accumulation of the blue-shifted M photointermediate at 403 nm, different from many H^+^-pumping rhodopsins like GPR and GR, which showed dominantly the accumulation of red-shifted photointermediates, N or O (Fig. S2A) ^21,22^. Additionally, when NmC was bound to NM-R1, an extra sharp peak at 484 nm appeared in the transient absorption spectra, indicating the absorption change of the bound carotenoids (Fig. S2A)^12^. Multiple-exponential analysis identified five photointermediates (Fig. S2B) and determined the photocycle of NM-R1 (Fig. 3I). There are two different initial state recovery from the M photointermediate equilibrated with N and O (M_3_/N/O) and from the M photointermediate (M_4_) directly, indicating the photocycle is blanched at M_2_. The absorption changes of NmC were observed in the late photointermediates after the M_1_ intermediate, implying that the large ternary structural changes of NM-R1 triggered by deprotonation of the retinal Schiff-base, affecting the structure of carotenoids (Fig. S2). Interestingly, the decay of the M_3_/N/O photointermediates was accelerated by the binding of NmC (Fig. 3I and S2A). These data indicated that the bound carotenoids to NM-R1 accelerated the photocycle turnover of the H^+^ pump.

Moreover, if photoexcitation was carried out at wavelengths shorter than 440 nm, the transient absorption signal was enhanced in the NM-R1–NmC complex compared to NM-R1 alone (Fig. S2C and S2D). This suggests that energy transfer from carotenoids to the retinal chromophore enhanced retinal isomerization, similar to observations in the Kin4B8 system (Fig. S2D)^12^. Based on the absorption of the sample and the ratio of the transient absorption change between NM-R1–NmC and NM-R1 alone, the EET efficiency was estimated to be approximately 11−15% (Fig. S2E). Since carotenoids absorbed photons, reducing the number of photons directly exciting the retinal chromophore, it is assumed that the ratios of transient absorption change were less than 1 at the range of 460−520 nm. These analyses showed that carotenoids bound to NM-R1 not only act as light-harvesting antennae but also photocycle-accelerating pigments.

To further investigate the effect of individual carotenoids, the transient absorption spectra of NM-R1–NsC complex and NM-R1–TsC complex were monitored. The transient absorption spectra of both complexes showed an extra peak, indicating that accompanied by the structural changes of NM-R1 during the photocycle, the structure of myxol and zeaxanthin also change (Fig. S3A and S3B). Multiple-exponential analysis showed that when myxol or zeaxanthin bound to NM-R1, the decay of the M_3_/N/O photointermediates was accelerated (Fig. S3B). Furthermore, the transient absorption signal was enhanced for both complexes, demonstrating that myxol and zeaxanthin function as light-harvesting antennae (EET efficiency: approximately 9% and 22%, respectively) (Fig. S3D). Therefore, both myxol and zeaxanthin function as photocycle-accelerating pigments and light-harvesting antennae.

In the NM-R3–myxol complex, the transient absorption measurements detected the accumulation of the red-shifted O-photointermediate at 620 nm, consistent with a previous study (Fig. S4A)^23^. While the shape of the ground-state bleach signal was a single peak at around 520 nm in the absence of myxol, it split into two peaks at 511 and 532 nm when myxol was bound to the protein (Fig. S4A). Additionally, when NM-R3 bound myxol, a band at ∼525 nm, reflecting myxol, appeared on the L_2_ and O_1_-photointermediates absorption spectra (Fig. S4A). These results indicated that accompanied by the structural changes of NM-R3 during the photocycle, the structure of myxol also changed, as in NM-R1. The fluorescence excitation spectrum (monitored at the retinal emission of 720 nm) of the NM-R3–myxol complexes exhibited three characteristic bands from myxol at 465, 500, and 535 nm, along with the 568 nm band of the retinal (Fig. 3J). The appearance of the myxol bands on the retinal excitation spectrum reflects that myxol transfers harvested light energy to the retinal leading to the formation of its excited singlet state (S_1_) and contributes to the emission from this state^24^. This observation demonstrates the direct energy transfer from the myxol to the retinal chromophore, proposing the possibility that myxol acts as an antenna to NM-R3. However, binding myxol to NM-R3 did not significantly enhance the transient absorption signal, indicating that light energy transferred from myxol did not promote retinal isomerization (Fig. S4C-E).

### Carotenoids bind to proteins in the different modes between NM-R1 and NM-R3

To examine the binding mode of carotenoids to rhodopsins, we used cryo-electron microscopy (cryo-EM) to determine the structures of rhodopsins–carotenoids complexes. We solved the structure of NM-R1–myxol complex and NM-R1–zeaxanthin complex at a resolution of 2.27 Å and 2.46 Å, respectively (Fig. 4A, 4B, and Fig. S5, S6, S7). NM-R1 forms a pentamer similar to that of GPR^25^. The multiple sequence alignment revealed NM-R1 has a motif sequence in the third helix, Asp-87, Thr-91, and Glu-98 (DTE motif) as specific amino acid residues (motif sequences) crucial for ion transport activity (Fig. S1). These correspond to the Asp-85, Thr-89, and Asp-96 (DTD motif) of HsBR. The structure of the proton translocation pathway and the function of NM-R1 as a light-driven proton pump indicates that NM-R1 possesses putative proton acceptor, Asp-87, and donor, Glu-98 (corresponding to Asp-85 and Asp-96 in HsBR, respectively) (Fig. S8) ^4^. In NM-R1, a myxol or a zeaxanthin molecule binds to each subunit and lies transversely against the outer surface of TM5 and TM6 at a 52° or 56° to the membrane normal, respectively (Fig. 4C and 4D). The myxol molecule binds close to a retinal molecule through the fenestration, and the distance between the centers and the ring moieties of two chromophores are 10.2 Å and 7.3 Å, respectively, and the angle between the axes of chromophores is 63°. A zeaxanthin molecule also binds to the same place: the distance between the centers and the ring moieties of two chromophores are 10.2 Å and 7.2 Å, respectively, and the angle between the axes of chromophores is 62°. Part of the retinal β-ionone ring is in van der Waals distance of the carotenoids β-ionone ring, and they are in contact with the Tyr-194 ring. These carotenoids β-ionone rings are immobilized by Trp-139, Leu-143, Met-195, Trp-201, and Tyr-202, amino acid residues in TM5 and TM6, and by the retinal β-ionone ring (Fig. 4E and 4F). The binding modes and overall structures of NM-R1 and carotenoids are largely similar to other rhodopsin-antenna complexes^10,12^.

**Figure 4.**
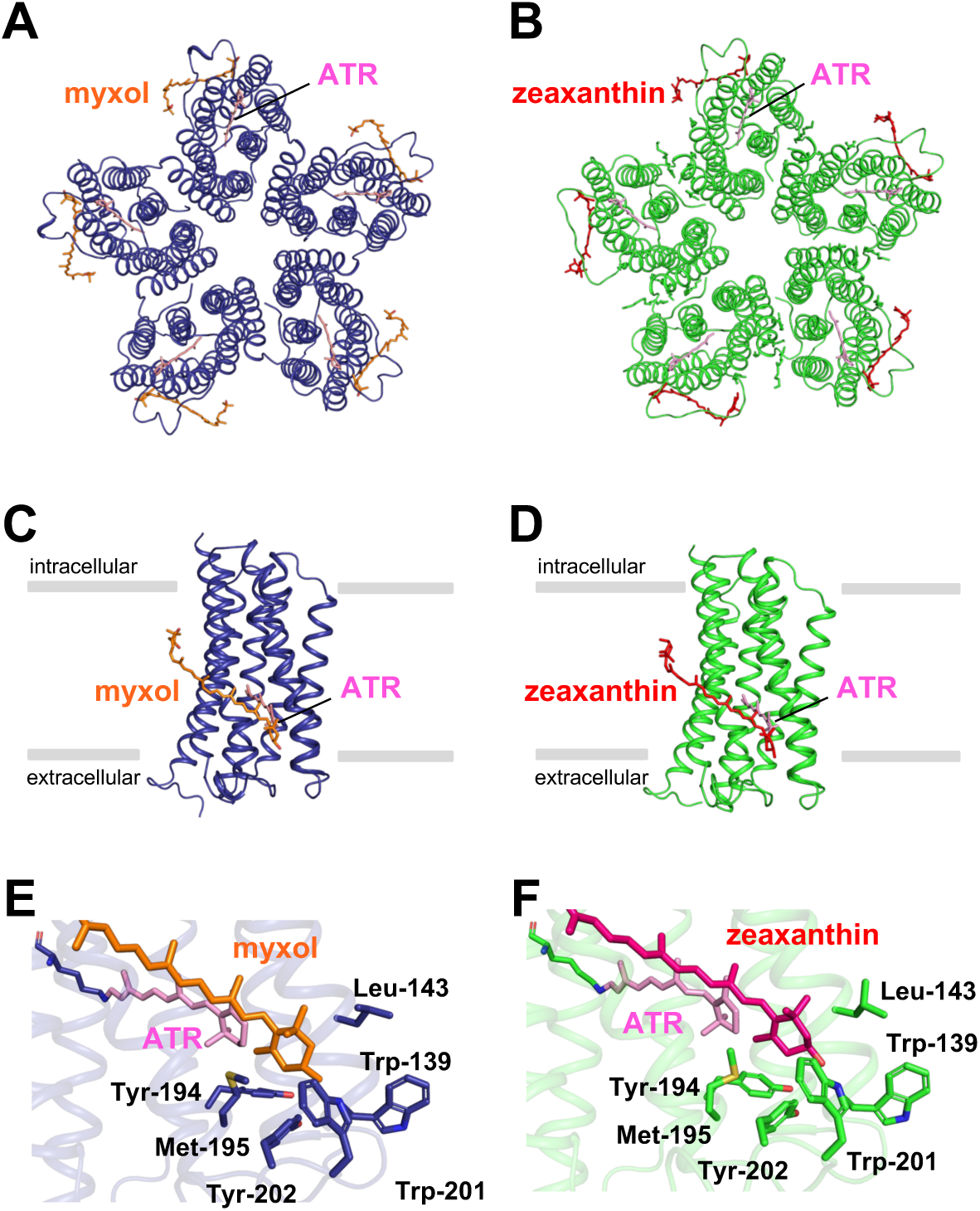
Cryo-EM structure of NM-R1. (A) Cryo-EM structure of pentameric NM-R1 structure showing from the extracellular side and the position of the all-*trans* retinal (ATR) and myxol. (B) Cryo-EM structure of pentameric NM-R1 structure showing from the extracellular side and the position of the retinal and zeaxanthin. (C) Side view of NM-R1 bound to myxol. (D) Side view of NM-R1 bound to zeaxanthin. (E) Positional relationship between the myxol, retinal, and surrounding residues of NM-R1. (F) Positional relationship between the zeaxanthin, retinal, and surrounding residues of NM-R1. Stick models of retinal, myxol, and zeaxanthin are shown in pink, orange, and red, respectively.

Next, we determined the cryo-EM structure of NM-R3–myxol complex at a resolution of 2.48 Å. NM-R3 forms a pentamer, with a myxol molecule binding to each subunit (Fig. 5A, S5, S9). However, unlike NM-R1, the myxol molecule lies longitudinal against the outer surface of TM5 and TM6 at a 12° to the membrane normal (Fig. 5B). A myxol molecule is located close to a retinal molecule, and the distance between the centers and the ring moieties of two chromophores are 11.7 Å and 7.9 Å, respectively. The angle between the axes of the chromophores is 56°. As well as NM-R1, part of the retinal β-ionone ring is in van der Waals distance of the myxol β-ionone ring through the fenestration, and they are in contact with the Tyr-208 ring. The binding site of the myxol β-ionone ring is formed by Met-154, Ala-158, Thr-161, Tyr-208, and Leu-209, amino acid residues in TM5 and TM6 (Fig. 5C). Toward the acyclic end containing a hydroxyl group, the carbon skeleton of myxol is immobilized by Phe-164, Thr-202, Val-165, Asn-168, Trp-169, Thr-173, Phe-176, and Lys-177, amino acid residues in TM5 and TM6 (Fig. 5D). Previously reported rhodopsin–antenna complexes conserve structural features and carotenoids lies transversely in common^10,12^. The structure of NM-R3–myxol complex greatly differs from them, suggesting that the binding mode of myxol to NM-R3 also differs from NM-R1 (Fig. 5E). To confirm this finding, we performed cryo-EM structural analysis of NM-R3 alone (Fig. S10). The results did not show the EM density of myxol observed in the NM-R3–myxol complex, supporting that the myxol arrangement observed in the NM-R3–myxol complex is correct.

**Figure 5.**
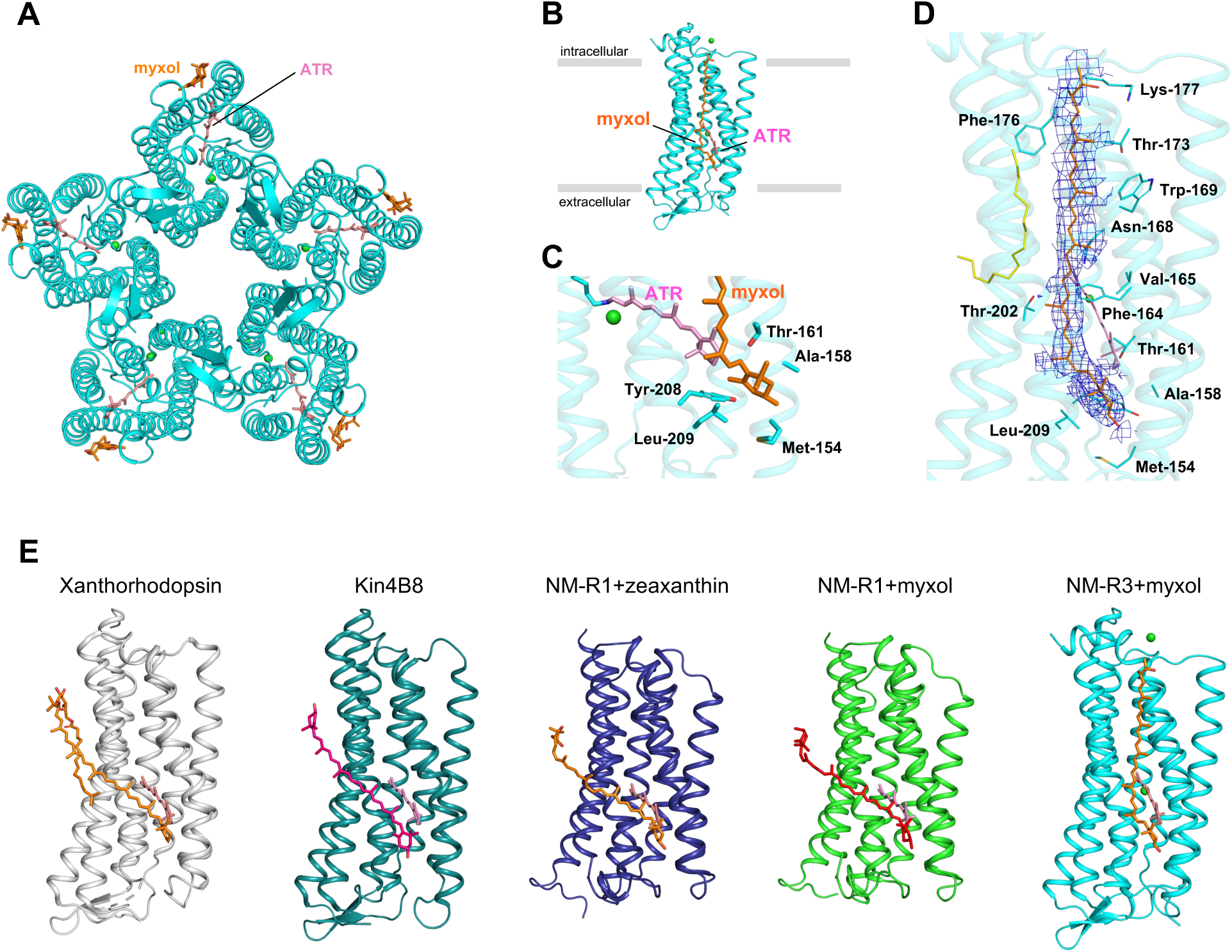
Cryo-EM structure of NM-R3. (A) Cryo-EM structure of pentameric NM-R3 structure showing from the extracellular side and the position of the all-*trans* retinal (ATR) and myxol. (B) Side view of NM-R3 bound to myxol. (C) Positional relationship between the myxol, retinal, and surrounding residues of NM-R3. (D) Myxol-binding site with the cryo-EM density of NM-R3. The extended carotenoid is tightly bound to the transmembrane surface of NM-R3. (E) Structure comparison between xanthorhodopsin (gray; PDB code 3DDL), Kin4B8 (deepteal; PDB code 8I2Z), NM-R1–myxol (deepblue), NM-R1–zeaxanthin (light green), and NM-R3–myxol (cyan). Stick models of retinal, myxol, and zeaxanthin are shown in pink, orange, and red, respectively. Chloride ions are shown in light green spheres.

### The characteristic residue for carotenoid binding is conserved in PRs of the marine *Bacteroidota* that are widely expressed in the photic zone

To assess the phylogenetic diversity of carotenoid-binding PRs in marine bacteria, a multiple sequence alignment and a phylogenetic tree was reconstructed using 1,009 PR sequences (Fig. 6A). PRs of bacteria belonging to the SAR11 clade, the most abundant group within the *Alphaproteobacteria* in the ocean, are known to carry either blue-absorbing PR (*λ*_max_: 490 nm, a color tuning residue at GPR position 105 is leucine) or green-absorbing PR (*λ*_max_: 525 nm, the color tuning residue is glutamine)^26^. A series of analyses revealed that both types of PRs of SAR11 have bulky residues (phenylalanine and tryptophane) at XR position 156, preventing carotenoid antennae binding (Fig. 6A). On the other hand, almost all PRs of *Bacteroidota* conserve the residue for green-light-absorbing residue (methionine as the color tuning residue)^27^ and a glycine residue at XR position 156, suggesting that they can bind carotenoids (Fig. 6A).

**Figure 6.**
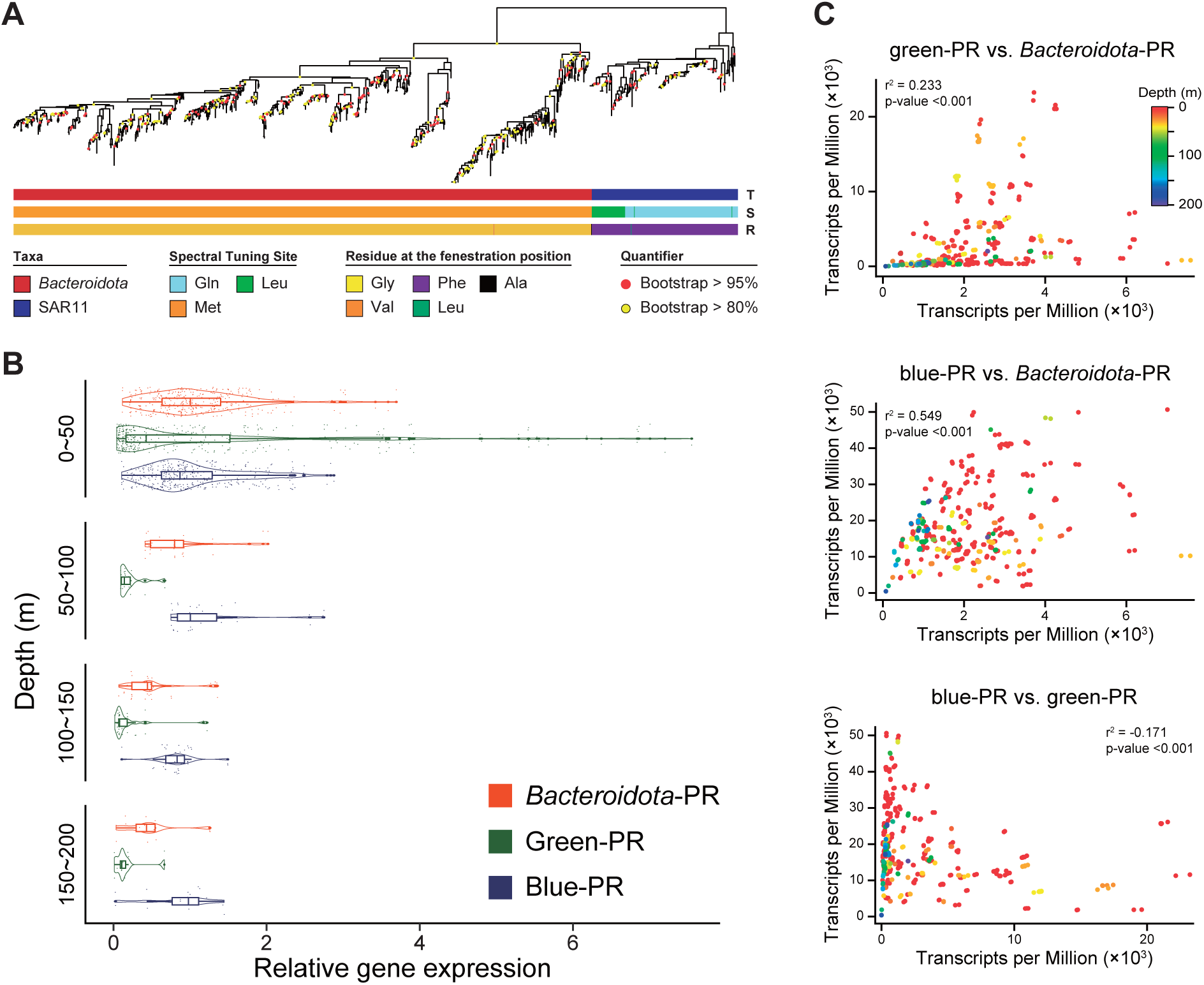
Environmental conditions favoring the use of blue-PRs, green-PRs, and antenna-containing PRs. (A) Phylogenetic tree of PR amino acid sequences. Associated taxa (top), spectral tuning site (middle; the leucine and the methionine residues: green-absorbing, the glutamine residue: blue-absorbing), and residue at the fenestration position (bottom) are color coded for each sequence. (B) *Bacteroidota*-PRs (left), SAR11 green-PRs (middle), and SAR11 blue-PRs (right) genes expression levels in the photic zone. The horizontal axis shows the relative gene expression levels to the average value of each type of PR gene expression at all depths. (C) Correlation analyses of green-PRs transcripts (y-axis) vs. *Bacteroidota*-PRs (x-axis) (top), blue-PRs transcripts (y-axis) vs. *Bacteroidota*-PRs (x-axis) (middle), and blue-PRs transcripts (y-axis) vs. green-PRs (x-axis) (bottom). Pearson correlation coefficient (r) values are 0.233, 0.549, and −0.171, respectively. All *P* values are less than 0.001. Circle color indicates the depth of each sample.

Because the carotenoid-binding ability of *Bacteroidota*-PRs expands their light absorption range, we analyzed metatranscriptomic data from the *Tara* Oceans project to assess whether PR gene expression varies with water depth. In general, blue light penetrates deeper than green light in the open ocean^12^, and the absorption wavelengths of PRs are closely linked to the depths at which organisms thrive. Metatranscriptomic analysis showed that SAR11 green-absorbing PR genes were mainly expressed in the shallow ocean layer (0–50 m), while SAR11 blue-absorbing PR genes were expressed over a broader depth range (0–200 m) (Fig. 6B). Interestingly, *Bacteroidota*-PR genes were also expressed across a wide vertical range, from 0 to 200 m (Fig. 6B). In the photic zone, *Bacteroidota*-PR genes expression patterns closely resembled those of both blue-absorbing PRs and green-absorbing PRs (vs. blue-PR is r^2^ = 0.549, *P* < 0.001; vs. green-PR is r^2^ = 0.233, *P* < 0.001). In contrast, blue-absorbing PRs and green-absorbing PRs showed a moderate inverse correlation (r^2^ = -0.171, *P* < 0.001) (Fig. 6C). These results suggest that the ability of *Bacteroidota*-PRs to bind carotenoid antennae, which expand their absorption range to blue-green spectral region, allows these bacteria to effectively capture light energy across a wide vertical range in the photic zone.

## Discussion

In this study, we investigated the carotenoid-binding abilities of three types of rhodopsins from marine bacteria and examined how these bound carotenoids affect rhodopsin properties. Spectroscopic analysis of the mixtures of intracellular carotenoids and heterologously expressed rhodopsins revealed that not only NM-R1 but also NM-R2 and NM-R3 can bind to carotenoids (Fig. 1D and 1E). Previously, carotenoid binding was limited to H^+^ transporting rhodopsins from the XLR and PR clades^12,19^. However, these findings demonstrate that other ion-transporting rhodopsins can also bind to carotenoids, broadening our understanding of rhodopsin–carotenoid interactions. While over 850 types of carotenoids exist in nature, only a few, such as salinixanthin, echinenone, canthaxanthin, nostoxanthin, lutein, and zeaxanthin, have been identified as light-harvesting antennae that enhance rhodopsin activity. These carotenoids are typically monocyclic 4-keto or dicyclic symmetric hydroxylated types^8,12,15,28–30^. In contrast, myxol, which was examined in this study, is a monocyclic, asymmetric hydroxylated carotenoid (see Fig. 2B). Monocyclic carotenoids with a γ-carotene backbone, such as myxol, are common among marine bacteria within the phylum *Bacteroidota*^31,32^. For example, marine bacterial isolates such as *Polaribacter irgensii* 23-P^T^ and *Psychroflexus torquis* ATCC 700755^T^ are known to carry rhodopsin genes and produce myxol^33–35^. These results suggest that marine *Bacteroidota* rhodopsins can form complexes with a wide variety of carotenoids, allowing flexibility in their light absorption.

Spectroscopic analyses revealed that carotenoids enhanced rhodopsin function in two distinct ways: by expanding the available light wavelengths and by accelerating the turnover of the photocycle. Absorption changes due to carotenoid binding were observed in the later photointermediates of NM-R1, and the carotenoids accelerate the later steps of the photocycle (such as the decay of M photointermediates, the accumulation and the decay of O photointermediates, and the recovery of the original state). This suggests that carotenoids may promote H^+^ uptake of NM-R1 (Fig. S11). Overall, the photocycle of NM-R1 was accelerated from 41 ± 10 ms (49%) or 130 ± 50 ms (51%) to about 32 ms by carotenoids (see Fig. 3I). Since light-harvesting antennae are the only reported mechanism to enhance rhodopsin reactivity to light, these photocycle-accelerating pigments represents an unprecedented type of interaction between carotenoids and rhodopsins. In NM-R3, structural analyses showed that myxol bound to each subunit of the pentamer and was located on the fenestration. These data, together with the excitation spectrum, indicate the possibility that myxol functions as an antenna, transferring the light energy to the retinal chromophore. As energy transfer from binding carotenoids to the retinal chromophore had been detected only in H^+^ pumping rhodopsins, this ClR–carotenoids complex indicates the diversity of rhodopsins–carotenoids interaction. However, since light energy transferred from myxol to the retinal chromophore did not promote the retinal isomerization, further study is needed to clarify what this transferred energy is used for, and what the biological significance of the light-harvesting antenna to ClR is.

The multiformity of rhodopsins–carotenoids interaction is also indicated by the fact that unlike NM-R1, NM-R3 is selectively bound to myxol. This selectivity can be attributed to the molecular structures of both the carotenoids and the rhodopsin. For instance, the cyclohexane ring structure of zeaxanthin likely acts as a steric barrier, obstructing interaction with the side chains of Thr-173, Phe-176, and Lys-177 and nearby lipid molecules, thereby blocking its binding to NM-R3 (Fig. S12). Additionally, unlike in other rhodopsin–carotenoid complexes, myxol binds vertically NM-R3. NM-R3 has a crevasse between TM5–TM6 where myxol can interact, but this crevasse is absent in NM-R1 due to a difference in the amino acid side chain, preventing carotenoid interaction along TM5–TM6 in NM-R1. Furthermore, the side chain of Thr-202 in NM-R3 locks the horizontal binding of myxol, whereas in NM-R1, the homologous amino acid is Ala-188, which presents no steric hindrance, allowing myxol to bind horizontally to NM-R1 (Fig. S13). Although the ecological role of the carotenoid selectivity remains unclear, these finding highlight that each type of rhodopsin in the same cell constructs a unique complex with carotenoids.

Phylogenetic analyses revealed that *Bacteroidota-*PRs form a distinct clade from PRs of *Pseudomonadota*^36^ including SAR11. SAR11 bacteria utilize PRs specialized for blue or green light absorption. In contrast, *Bacteroidota* have conserved a unique carotenoid-binding PR lineage, allowing them to utilize intracellular carotenoids as light-harvesting antennae and absorb a broader spectrum of light. Moreover, metatranscriptomic analysis showed that *Bacteroidota*-PRs were expressed across a wide vertical range in the photic zone, underscoring the role of carotenoids in enhancing the *Bacteroidota*-PR mediated phototrophy under various light conditions, which support their growth and survival across varying depths^36–38^.

It was thought that the only mechanism to enhance rhodopsin-mediated phototrophy was the light-harvesting antenna in H^+^-pumping rhodopsins. This study showed that carotenoids bound to H^+^-pumping rhodopsins not only work as antennae, but also as photocycle-accelerating pigments. In addition, a Cl^−^-pumping rhodopsin containing a carotenoid antenna was discovered. Together, these findings expand our understanding of how rhodopsin-mediated phototrophy adapts in the marine environment, revealing diverse strategies that marine bacteria use to capture and utilize light energy.

## Limitation of this study

While our findings provide unprecedented insights into the mechanisms by which carotenoids enhance rhodopsin functions, several challenges about the detailed characteristics of rhodopsin–carotenoid complexes remain. First, the precise molecular mechanism underlying photocycle accelerating is still unclear, despite our observation of this phenomenon. Second, the dynamics of light energy transfer from the myxol to the retinal and how it affects NM-R3 are not fully understood. Finally, we did not quantify the potential increase in ion transport rates due to photocycle-accelerating pigments. Further studies will be necessary to address these questions and deepen our understanding of the molecular interactions between carotenoids and rhodopsins.

## Resource availability

### Lead contact and Materials availability

This study did not generate new unique reagents. Further information and requests for resources and reagents should be directed to and will be fulfilled by the lead contact, Susumu Yoshizawa.

## Data availability

All data are available in the main text or the supplementary information. The density map and structure coordinate of the cryo-EM structure of NM-R1–myxol complex, NM-R1– zeaxanthin complex, NM-R3–myxol complex, and NM-R3 have been deposited in the Electron Microscopy Data Bank and the Protein Data Bank with accession numbers EMD-61686, 9JOV, EMD-61685, 9JOU, EMD-61687, 9JOW, EMD-61688, and 9JOX, respectively.

## Acknowledgements

We sincerely thank Prof. Oded Béjà at Technion-Israel Institute of Technology for his comments on our experiments and paper. This work was supported by MEXT Advancement of Technologies for Utilization Big Data of Marine Life (grant JPMXD1521474594), JSPS KAKENHI Grants-in-Aid (grants 22KJ1110 to T.F., JP23H04404 to K.I., and 22H00557 to S.Y.), JST CREST (grant JPMJCR22N2 to K.I.), Research Support Project for Life Science and Drug Discovery (Basis for Supporting Innovative Drug Discovery and Life Science Research (BINDS)) from AMED (grant JP23ama121013 to M.S.), and MEXT Promotion of Development of a Joint Usage/ Research System Project: Coalition of Universities for Research Excellence Program (CURE) (grant JPMXP1323015482 to K.I.). All cryo-EM data in this study were collected at the cryo-EM facility of the RIKEN Center for Biosystems Dynamics Research (Yokohama, Japan).

## Author contributions

T.F. and S.Y. designed the research; T.F., M.H.-T., Y.T. and S.Y. measured absorption and emission spectra of NM-R1 and NM-R3, culturing marine *Bacteroidota*. T.H., T.U.-K., K.H., and M.S. performed structural analysis. Y.N., K.T., K.M., and S.N. performed bioinformatics. K.I. performed CD spectra measurements and laser-flash photolysis. S.T. and T.M. identified carotenoid pigments. T.F. and S.Y. wrote the paper with input from all authors.

## Supplemental information

Figures S1–S13 and Table S1–S2.

## Star★Methods

Detailed methods are provided in the online version of this paper and include the following:

### Phylogenetic analysis

For the phylogenetic analysis of rhodopsins, amino acid sequences of 49 microbial rhodopsins were collected from the National Center for Biotechnology Information and aligned using MAFFT version 7.515 with EINSI strategy^39^. The phylogenetic tree was inferred using RAxML version 8.2.12 with the PROTGAMMALGF model using 1,000 rapid bootstrap searches^40^. Model selection was performed with the ProteinModelSelection.pl script in the RAxML package. The phylogenetic trees were visualized with Interactive tree of Life version 6.8.1^41^.

### Metagenomic datasets

For marine Bacteroidets, we used the 1,560 representative metagenome-assembled genomes (MAGs) of *Bacteroidota* species, which were reconstructed in a previous study^42^. For SAR11, the genomes from multiple studies, including MAGs, genomes of isolated strains, and Single Amplified Genomes (SAGs), were collected according to the previous studies^43–62^, because producing MAGs from predominant marine prokaryoplankton such as SAR11 is difficult. Sequence reads of the 509 meta-transcriptomes obtained in the *Tara* Ocean samples were mapped onto these genomes^50,53,63,64^. We downloaded paired-end sequence data from NCBI SRA by SRA toolkit (https://github.com/ncbi/sra-tools), and then quality control was performed by using fastp with options -q 20 -n 10 -l 60^65^. The quality-controlled reads were mapped via bwa-mem2^66^. Transcripts per million (TPM) were calculated using a script from a previous study (https://github.com/yosuken/CountMappedReads2)^67^.

Gene prediction from these genomes was performed by the prokka pipeline^68^. To extract PR, genes assigned as COG5524 (Bacteriorhodopsin) by COGclassifier (https://github.com/moshi4/COGclassifier)^69^ and PF01036 (Bac_rhodopsin) by HMMER (sequence and domain score cutoff: 24.6) were collected^70^. Further, orthologous genes that were clustered in the same orthologue with these genes by SonicParanoid2 were also collected^71^. Too short sequences (length <150 AA) were discarded.

To infer the phylogenetic tree of the PR genes, the amino acid sequences of the CDSs were aligned by MAFFT version 7.212 with global alignment mode^72^. The sequences lacking the residues associated with the light absorption spectrum and carotenoid antennae were discarded after alignment. The alignments were curated using trimAl version 1.4.1^73^. The maximum-likelihood trees were inferred using IQ-TREE version 2.2.0 with option -m MFP and -bb 1000^74–76^. Tree files were visualized using ggtree in R^77,78^.

### Rhodopsin expression

Codon-optimized DNA fragments for *Escherichia coli* encoding NM-R1 (GenBank: BAO54171.1), NM-R2 (GenBank: BAO54106.1), and NM-R3 (GenBank: BAO55276.1) were chemically synthesized (Eurofins Genomics) and inserted into the pET21a(+) plasmid vector (Novagen). The plasmids were transformed into *E.coli* strain C41 (DE3) (Lucigen) cells, and colonies were inoculated to Luria-Bertani (LB) medium supplemented with 100 μg mL^−1^ ampicillin. Before protein expression, transformants were cultured at 37 °C in 2× YT medium (NaCl 5 g L^−1^, Bacto tryptone 16 g L^−1^, and Bacto yeast extract 10 g L^−1^, pH 7.0) with 100 μg mL^−1^ ampicillin until the OD_660_ reached 0.3. Protein expression was then induced at 37 °C for 4 h in the dark by adding 0.1 mM isopropyl β-D-1-thiogalactopyranoside (IPTG: Sigma-Aldrich) and 10 μM all-*trans* retinal (Sigma-Aldrich). Rhodopsin-expressing cells were collected by centrifugation at 4,400 × *g* for 3 min at 4 °C (MX-305, Tomy Seiko) and then washed twice in 100 mM NaCl.

### Protein purification and absorbance measurements

Rhodopsin-expressing *E.coli* cells were resuspended in 7 mL of buffer containing 50 mM Tris–HCl pH 8.0 and 500 mM NaCl, and disrupted by sonication (Branson SFX 250 Digital Sonifier, Branson Ultrasonics) on ice-cold water for 5 min. Crude membranes were collected by ultracentrifugation at 106,800 × *g* for 30 min at 4 °C (Optima XPN-90 Ultracentrifuge with a SW 32Ti rotor, Beckman Coulter) and solubilized with 1% (w/v) *n*-dodecyl-β-D-maltoside (DDM; Dojindo Lab). The solubilized rhodopsin proteins were obtained by ultracentrifugation at 106,800 × *g* for 30 min at 4 °C (Optima XPN-90 Ultracentrifuge with a SW 32Ti rotor, Beckman Coulter) and purified by a HisTrap FF Ni-NTA affinity chromatography column (GE Healthcare) at room temperature (approximately 25 °C). The purified sample was concentrated, and its buffer was replaced with a new buffer containing 25 mM MOPs (pH 7.2), 500 mM NaCl, and 0.1% DDM using Amicon 30-kDa cut-off (UFC803024, Millipore) by centrifugation at 5,000 × *g* for 20 min at 4 °C (MX-305, Tomy Seiko). UV-Vis absorption spectral measurements of rhodopsins were taken with a UV-2600 spectrophotometer (Shimadzu) at room temperature (approximately 25 °C).

### Chromophores extraction from marine bacterial cells

*Nonlabens marinus* S1-08^T^, *Tenacibaculum* sp. SG-28, and *N. spongiae* JCM 13191^T^ cells were cultured in 500 mL half-strength ZoBell’s 2216E medium^79^ under light conditions at 25 °C until the OD_600_ reached 0.1. Cultured cells were collected by centrifugation at 4,400 × *g* for 10 min at 20 °C (MX-305, Tomy Seiko). After washing cells twice in 100 mM NaCl, intracellular pigments were extracted with 30 mL of acetone/methanol (7:3, v/v). Air-dried pigments were resuspended in 30 mL of the buffer containing 25 mM MOPs (pH 7.2), 500 mM NaCl and 0.1% DDM, and filtered through a 0.22 μm pore size filter (Advantech) to remove insoluble pigments. The filtered solution was concentrated, using Amicon 30-kDa cut-off (UFC903024, Millipore) by centrifugation at 5,000 × *g* for 20 min at 4 °C (MX-305, Tomy Seiko). UV-Vis absorption spectral measurements of chromophores were taken in the same way as purified proteins.

### Binding of chromophore extracted from cells to rhodopsin proteins

Purified rhodopsin proteins and NmC were mixed in a ratio of 450 μL (A_530_ of 0.5) to 600 μL (A_520_ of 1.5) and incubated overnight with gentle rotation at 4 °C in the dark. The mixed samples were purified using a HisTrap FF Ni-NTA affinity chromatography column (Takara Bio) at 4 °C. The concentration and replacement of the buffer were carried out in the same way as protein purification.

### Chromatographic analysis of carotenoids extracted from N. marinus *S1-08^T^* cells and carotenoids bound to rhodopsins

*Nonlabens marinus* S1-08^T^ cells were cultured and collected as described above. Intracellular pigments were extracted with 7 mL of acetone/methanol (7:3, v/v). Cells are disrupted by sonication (Branson SFX 250 Digital Sonifier, Branson Ultrasonics) on ice-cold water for 15 s, and cell debris was removed by centrifugation at 10,000 × *g* for 10 min at 4 °C (MX-305, Tomy Seiko). Purified rhodopsin–carotenoid complexes (A_520_ of 1.0) of 1 mL were concentrated to 100 μL using Amicon 30-kDa cut-off (UFC903024, Millipore) by centrifugation at 5,000 × *g* for 20 min at 4 °C (MX-305, Tomy Seiko) and were added to 500 μL of methanol, and carotenoid pigments bound to rhodopsin were extracted. These solutions were evaporated at 25 °C and air-dried pigments were resuspended in 100 μL methanol and settled at –80 °C for 3 h. Solutions were filtered through a 0.22 μm pore size spin-filter (Ultrafree, Millipore) by centrifugation at 14,200 × *g* for 5 min at 4 °C (MX-305, Tomy Seiko). For HPLC analysis, samples were prepared to mix solutions and ultrapure water (8:2, v/v).

Carotenoid pigments were detected in a HPLC (Prominence-i LC-2030, Shimadzu), equipped with a PDA detector, using a Kinetex 5 μm EVO C18 100A column (Phenomenex) at a flow rate of 2 mL min^−1^. The solvent system was a mixture of H_2_O/methanol (2:8, v/v, solvent A) and methanol (solvent B). The eluent was linear gradient for 0–6 min, 100% A to 50% A:50% B; 6–40 min, 50% A:50% B to 40% A:60% B. The column temperature was kept at 60 °C, and the injection volume was 20 μL. LabSolutions software was used for data processing.

### Carotenoid identification

Pigments were extracted with acetone/methanol (7:3, v/v). Carotenoids were separated from phospholipids using a DEAE-Toyopearl 650 M column chromatography eluted with hexane/acetone (1:1, v/v). Next, two major carotenoids were collected from silica gel TLC developed with dichloromethane/ethyl acetate (4:1, v/v) and then the purified carotenoids were collected from C18-HPLC eluted with methanol. Molecular mass of the peak 1 component was 607.4125 [MNa]^+^ (calculated 607.4127 for C_40_H_56_O_3_Na) and that of the peak 2 was 569.4332 [MH]^+^ (calculated 569.4359 for C_40_H_57_O_2_). ^1^H-NMR were measured in CDCl_3_, and CD were measured in diethyl ether/*i*-pentane/ethanol (5:5:2, by vol.).

### Hydroxylamine bleaching

Purified NM-R1 and NmC were mixed in a ratio of 500 μL (A_525_ of 0.3) to 500 μL (A_511_ of 1.0). Purified NM-R1 and TsC or NsC were mixed in a ratio of 450 μL (A_525_ of 0.25) to 600 μL (A_460_ or A_483_ of 0.8, respectively).

Purified NM-R3 and chromophores extracted from *Nonlabens marinus* S1-08^T^ cells were mixed in a ratio of 500 μL (A_530_ of 0.4) to 500 μL (A_520_ of 1.2). These mixtures were incubated overnight with gentle rotation at 4 °C in the dark. The mixture of NM-R1 and NM-R3 with chromophores were bleached with 100 and 250 mM hydroxylamine, respectively. To promote the reaction, NM-R3 and NmC mixture was illuminated with light from a 300 W Xenon lamp (MAX-303, Asahi Spectra) through a bandpass filter (λ = 520 ± 10 nm; MX0520, Asahi Spectra). The absorption changes were monitored using a UV-2600 spectrophotometer (Shimadzu) at room temperature (approximately 25 °C).

### Purification of rhodopsin-nanodiscs

The sequence encoding NM-R1 was cloned into pCR2.1-TOPO (Thermo Fisher Scientific Life Sciences) with C-terminal tobacco etch virus (TEV) protease digestion site, Strept-tag II, and histidine tags, and was expressed in *E. coli* Lemo21 (DE3) cells (New England Biolabs). *E. coli* cells containing NM-R1 expression vector were grown in LB medium supplemented with 50 µg mL^−1^ ampicillin at 120 rpm at 37 °C. When the OD_600_ reached 1.0, expression was induced using a 1 mM final concentration of IPTG and 40 µM all-*trans* retinal. The induced culture was grown at 220 rpm for more than 4 h at 37 °C.

NM-R1-expressing *E. coli* cultures were centrifuged at 5,000 × *g* for 10 min at 4 °C and the pellet was resuspended in a buffer containing 50 mM HEPES pH 7.4, 150 mM NaCl. The sample was disrupted by using a French press for three passes at 1500 psi. Then, the sample was centrifuged at 5,000 × *g* for 10 min at 4 °C to pellet undisrupted cells or large cell debris. Membranes were collected by centrifuging the sample at 100,000 × *g* for 0.5 h at 4 °C and resuspended in a buffer containing 25 mM MOPs pH 7.2, 500 mM NaCl and 1% DDM final concentration. The sample was incubated for more than 2 h at 4 °C with gentle rotation and a second centrifugation at 100,000 × *g* for 0.5 h at 4 °C was performed. The supernatant was then incubated with Strept-tactin sepharose (Cytiva) for 1 h. Beads were washed on a gravity column using a buffer containing 25 mM MOPs pH 7.2, 500 mM NaCl and 0.04% DDM. Protein was eluted from the column using a buffer containing 25 mM MOPs pH 7.2, 500 mM NaCl, 0.04% DDM and 10 mM desthiobiotin.

Cell-free protein synthesis of NM-R3 was performed essentially according to the previously reported protocols^80,81^. After solubilization in 1.0% DDM, the supernatant containing NM-R3 was affinity purified on Ni-NTA resin (QIAGEN) with buffer (25 mM MOPs pH 7.2, 500 mM NaCl and 0.04% DDM, and 500 mM imidazole). The eluted protein of NM-R1 and NM-R3 were concentrated using Amicon 50-kDa cut-off (Millipore) and loaded onto a HiLoad Superdex 200 column equilibrated with 25 mM MOPs pH 7.2, 500 mM NaCl and 0.04% DDM. The collected fractions were concentrated using Amicon 50-kDa cut-off (Millipore).

Purified NM-R1 proteins and myxol or zeaxanthin were mixed in a ratio of 450 μL (A_530_ of 0.5) to 600 μL (A_520_ of 1.8) and incubated overnight with gentle rotation at 4 °C in the dark. The mixed samples were purified using a Strept-tactin sepharose (Cytiva) at 4 °C and subjected to TEV protease digestion. The cleaved tag and TEV protease were removed by passage through the Ni-NTA column, and the NM-R1-myxol or NM-R1-zeaxanthin proteins were recovered from the flowthrough fraction.

Purified NM-R3 proteins and myxol were mixed in a ratio of 450 μL (A_530_ of 0.8) to 600 μL (A_520_ of 1.8) and incubated overnight with gentle rotation at 4 °C in the dark. The mixed samples were purified using a Ni-NTA resin (QIAGEN) at 4 °C. The eluate was dialyzed against buffer (25 mM MOPs pH 7.2, 500 mM NaCl and 0.04% DDM). The affinity-tag was cleaved with TEV protease. The cleaved tag and TEV protease were removed by passage through the Ni-NTA column, and the NM-R3–myxol protein was recovered from the flowthrough fraction.

For the cryo-EM grid preparation of the NM-R1–myxol, NM-R1–zeaxanthin, NM-R3–myxol and NM-R3, the protein was reconstituted in nanodiscs, MSP1E3D1. Each rhodopsin, MSP1E3D1, and eggPC were mixed at a molar ratio of 3.6:2:50, respectively, and incubated at 4 °C for 1 h. Detergents were removed by adding Bio-Beads SM2 (Bio-Rad) and were incubated at 4 °C overnight. The Bio-Beads were then removed, and the samples were purified by size-exclusion chromatography on a Superdex 200 Increase 10/300 GL column (Cytiva), equilibrated with buffer containing 25 mM MOPs pH 7.2, 500 mM NaCl. The peak fractions of the protein were collected and concentrated to 4.0 mg mL^−1^, using a centrifugal filter unit (50 kDa molecular weight cut-off; Millipore). Of proteins, 4 µL was loaded onto glow-discharged Quantifoil R1.2/1.3 300 mesh holey Au grid using a Vitrobot Mark IV instrument (Thermo Fisher Scientific) with a blotting force of -3 for 3 s at 100% humidity and 4 °C.

### Circular dichroism spectroscopic measurement

Circular dichroism (CD) spectroscopic measurements were performed using a J-725 spectrometer (Jasco) at 23 °C. The rhodopsin-nanodisc sample solution, prepared in 25 mM MOPs–NaOH PH 7.2, 500 mM NaCl, was placed in a quartz cell cuvette with a 1 cm path length. Optical density of the samples was adjusted to ∼0.8 at the absorption maximum wavelengths. CD spectra were recorded with a bandwidth resolution of 1 nm, at 1 nm intervals, and a scan speed of 100 nm min^−1^.

### Fluorescence spectroscopic measurements

For fluorescence measurements, the buffer of rhodopsins-nanodiscs samples was replaced with a new buffer containing 25 mM MOPs pH 7.2, 100 mM Na_2_SO_4_, and 10 mM NaCl using Amicon 30-kDa cut-off (UFC903024, Millipore) by centrifugation at 5,000 × *g* for 20 min at 4 °C (MX-305, Tomy Seiko). The samples used for measurements for the rhodpsins were A_550_ of 0.148, and rhodopsins–myxol complexes were A_528_ of 0.12. Fluorescence emission and excitation spectra were measured on the RF-6000 spectrofluorometer (Shimadzu) at room temperature (approximately 25 °C). The slit width of both the emission and the excitation channel was 5 nm. Scattered excitation light was blocked by a long-pass filter (more than 510 ± 5 nm; LV0510, Asahi Spectra). The emission and excitation spectra were sampled at 530 nm and 720 nm, respectively. The following equation relating the fluorescence excitation spectral profile to the corresponding absorption spectral profile used to correct the inner filter effect: *F*_ideal_ = *F*_obs_10^(A_EX_ + A_Em_)/2^, where *F*_ideal_ is the ideal fluorescence intensity absence of the inner filter effect, *F*_obs_ is the measured fluorescence intensity, and A_Ex_ and A_Em_ are the absorbance at the excitation and emission wavelengths, respectively^82^.

### Laser-flash photolysis

For the laser flash photolysis spectroscopy, rhodopsin-nanodisc sample solution, prepared in 25 mM MOPs-NaOH PH 7.2, 500 mM NaCl, was used. The optical density of the retinal in rhodopsins was adjusted to ∼0.4–0.5 (protein concentration ∼0.2–0.25 mg mL^−1^) at the absorption maximum wavelengths. The laser flash photolysis measurement was conducted as previously described^83^. The nano-second second harmonics of Nd–YAG laser (*λ* = 532 nm, INDI40, Spectra-Physics) was used for the excitation of rhodopsins to determine the transient absorption spectra and the time courses of the transient absorption change at specific probe wavelengths. The transient absorption spectra were obtained by monitoring the intensity change of white light from a Xe-arc lamp (L9289-01, Hamamatsu Photonics), passing through the sample, with an ICCD linear array detector (C8808-01, Hamamatsu). To increase the signal-to-noise (S/N) ratio, 60–100 spectra were averaged, and the singular-value-decomposition (SVD) analysis was applied. To measure the time courses of transient absorption change at specific probe wavelengths, the output of a Xe-arc lamp (L9289-01, Hamamatsu Photonics) was monochromated by monochromators (S-10, SOMA OPTICS) and the change in the intensity after the photo-excitation was monitored with a photomultiplier tube (R10699, Hamamatsu Photonics). To increase S/N ratio, 100–200 signals were averaged.

The transient absorption change enhancement indicating the energy transfer from carotenoids to retinal was investigated by measuring the transient absorption change with sufficiently low (0.38 mJ cm^−2^) excitation pulse energy (Fig. S2C) to keep the linearity between absorbed photon number and signal intensity. Nano-second pulses from an optical parametric oscillator (basiScan, Spectra-Physics) pumped by the third harmonics of Nd–YAG laser (*λ* = 355 nm, INDI40, Spectra-Physics) were used for the excitation of rhodopsins at different wavelengths.

### Cryo-EM single partice analysis of the rhodopsin-antenna pigment complex

Cryo-EM imaging was performed on a Titan Krios G4 (Thermo Fischer Scientific) operated at 300 kV, equipped with a Gatan Quantum-LS Energy Filter (slit width 15 eV) and a Gatan K3 direct electron detector at a nominal magnification of 105,000× in electron-counting mode, corresponding to a pixel size of 0.83 Å per pixel. Stack movie of NM-R1–myxol, NM-R1–zeaxanthin, NM-R3–myxol and NM-R3 was recorded in an accumulated exposure of 60.4, 54.0, 60.4 and 57.5 e^−^ Å^−2^, respectively. These data were automatically acquired by the image-shift method using the EPU software with a defocus range of −0.8 to −2.0 μm. In the dataset, 5,000, 9,002, 5,314, and 5,000 movie stacks of NM-R1–myxol, NM-R1–zeaxanthin, NM-R3–myxol and NM-R3 were acquired, respectively. All image processing was performed with CryoSPARC^84,85^. A total of 4,709,010, 8,804,012, 5,747,079, and 5,620,703 of NM-R1–myxol, NM-R1–zeaxanthin, NM-R3–myxol and NM-R3 particles were picked from the micrographs and extracted at a pixel size of 200 Å, respectively. These particles were subjected to several rounds of 2D classifications. The selected 897,953, 1,814,697, 791,219, and 2,081,189 particles of NM-R1–myxol, NM-R1–zeaxanthin, NM-R3–myxol and NM-R3 were subjected to Non-Uniform refinement^86^ of the map improved its global resolution to 2.27, 2.46, 2.48, and 2.50 Å, according to the Fourier shell correlation (FSC) = 0.143^87^ criterion, respectively. The initial model of the NM-R1 and NM-R3 complex were built using Coot^88^ to fit models of the green-light absorbing proteorhodopsin (PDB code 7B03^89^) and the crystal structure of NM-R3 (PDB code 5B2N^81^) into the cryo-EM map, respectively. The high-resolution cryo-EM map allowed side-chain assignments for both NM-R1 and NM-R3 complex (Fig. S6). The entire structure was further manually adjusted and refined using PHENIX^90^ with phenix.real_space refine and servalcat^91^. The data collection, processing, refinement and validation statistics of the NM-R1–myxol, NM-R1–zeaxanthin, NM-R3– myxol and NM-R3 structure are listed in Supplementary Table S2.

