## supplemental Files for "Carotenoid pigments enhance rhodopsin-mediated phototrophy by light-harvesting and photocycle-accelerating"

### Supplemental information

Figures S1–S13 and Table S1–S2.

Figure S1 | The amino acid sequence alignment of important residues for the function of microbial rhodopsins.

Figure S2 | The photocycle of NM-R1 and NM-R1–NmC complex.

Figure S3 | The photocycle of NM-R1–NsC and NM-R1–TsC complex.

Figure S4 | The photocycle of NM-R3 and NM-R3–NmC complex.

Figure S5 | Cryo-EM density map shown in four proteins viewed from the extracellular side of the membrane and viewed from the side

Figure S6 | Structural feature of NM-R1 bound the myxol.

Figure S7 | Structural feature of NM-R1 bound the zeaxanthin.

Figure S8 | Structure of the proton translocation pathway of NM-R1 bound the myxol or zeaxanthin.

Figure S9 | Structural feature of NM-R3 bound the myxol.

Figure S10 | Structural feature of NM-R3.

Figure S11 | Comparison of time course of the transient absorption change of NM-R1 without carotenoids, with NmC, TsC, and NsC.

Figure S12 | Close up view of the myxol binding site of NM-R3.

Figure S13 | Myxol binding site of NM-R1 or NM-R3.

Table S1 | $^{1}$H-NMR data of myxol and zeaxanthin in $CDCl_3$.

Table S2 | Cryo-EM data collection, refinement, and validation statistics.

| No. in BR | | 82 | 83 | 85 | 86 | 89 | 96 | 138 | 182 | 212 | 216 |
|---|---|---|---|---|---|---|---|---|---|---|---|
| No. in XR | | 93 | 94 | 96 | 97 | 100 | 107 | 156 | 200 | 236 | 240 |
| | HsBR | R | Y | D | W | T | D | W | W | D | K |
| | XR | R | Y | D | W | T | E | G | W | D | K |
| PR clade | BPR | R | Y | D | W | T | E | F | W | D | K |
| | GPR | R | Y | D | W | T | E | F | W | D | K |
| | EINA29G6 | R | Y | D | W | T | E | G | W | D | K |
| | KR1 | R | Y | D | W | T | E | G | W | D | K |
| | TsPR | R | Y | D | W | T | E | G | W | D | K |
| | NM-R1 | R | Y | D | W | T | E | G | W | D | K |
| ClR clade | NM-R3 | R | Y | N | W | T | Q | G | W | D | K |
| | NsClR | R | Y | N | W | T | Q | G | W | D | K |
| | FR | R | Y | N | W | T | Q | G | W | D | K |
| | EryClR | R | Y | N | W | T | Q | G | W | D | K |
| | QaClR | R | Y | N | W | T | Q | G | W | D | K |
| NaR clade | IaNaR | R | Y | N | W | D | Q | G | W | D | K |
| | NdNaR | R | Y | N | W | D | Q | G | W | D | K |
| | NM-R2 | R | Y | N | W | D | Q | G | W | D | K |
| | GLR | R | Y | N | W | D | Q | G | W | D | K |
| | KR2 | R | Y | N | W | D | Q | G | W | D | K |
| | DoNaR | R | Y | N | W | D | Q | G | W | D | K |

**Figure S1 |** The amino acid sequence alignment of important residues for the function of microbial rhodopsins (orange, acidic; blue, basic; black, aromatic; green, –OH bearing; magenta, asparagine and glutamine residues; red, glycine), related to Figure 1. Abbreviations: HsBR, bacteriorhodopsin from *Halobacterium salinarum*; XR, xanthorhodopsin from *Salinibacter ruber* M31$^{T}$; BPR, blue-absorbing proteorhodopsin; GPR, green-absorbing proteorhodopsin; EINA29G6, KR1, TsPR, and NM-R1, PR from uncultured freshwater flavobacterium, *Krokinobacter eikastus* NBRC 100814$^{T}$, *Tenacibaculum* sp. SG-28, *Nonlabens marinus* S1-08$^{T}$, respectively; NM-R3, NsClR, FR, EryClR, and QaClR, ClR from *N. marinus* S1-08$^{T}$, *Nonlabens spongiae* UST030701-156$^{T}$, *Fulvimarina pelagi*, *Erythrobacter* sp. QSSC1-22B, and *Qipengyuania aestuarii*, respectively; IaNaR, NdNaR, NM-R2, GLR, KR2, and DoNaR, NaR from *Indibacter alkaliphilus*, *Nonlabens dokdonensis*, *N. marinus* S1-08$^{T}$, *Gillisia limnaea*, *K. eikastus* NBRC 100814$^{T}$, and *Dokdonia* sp. PRO95, respectively.

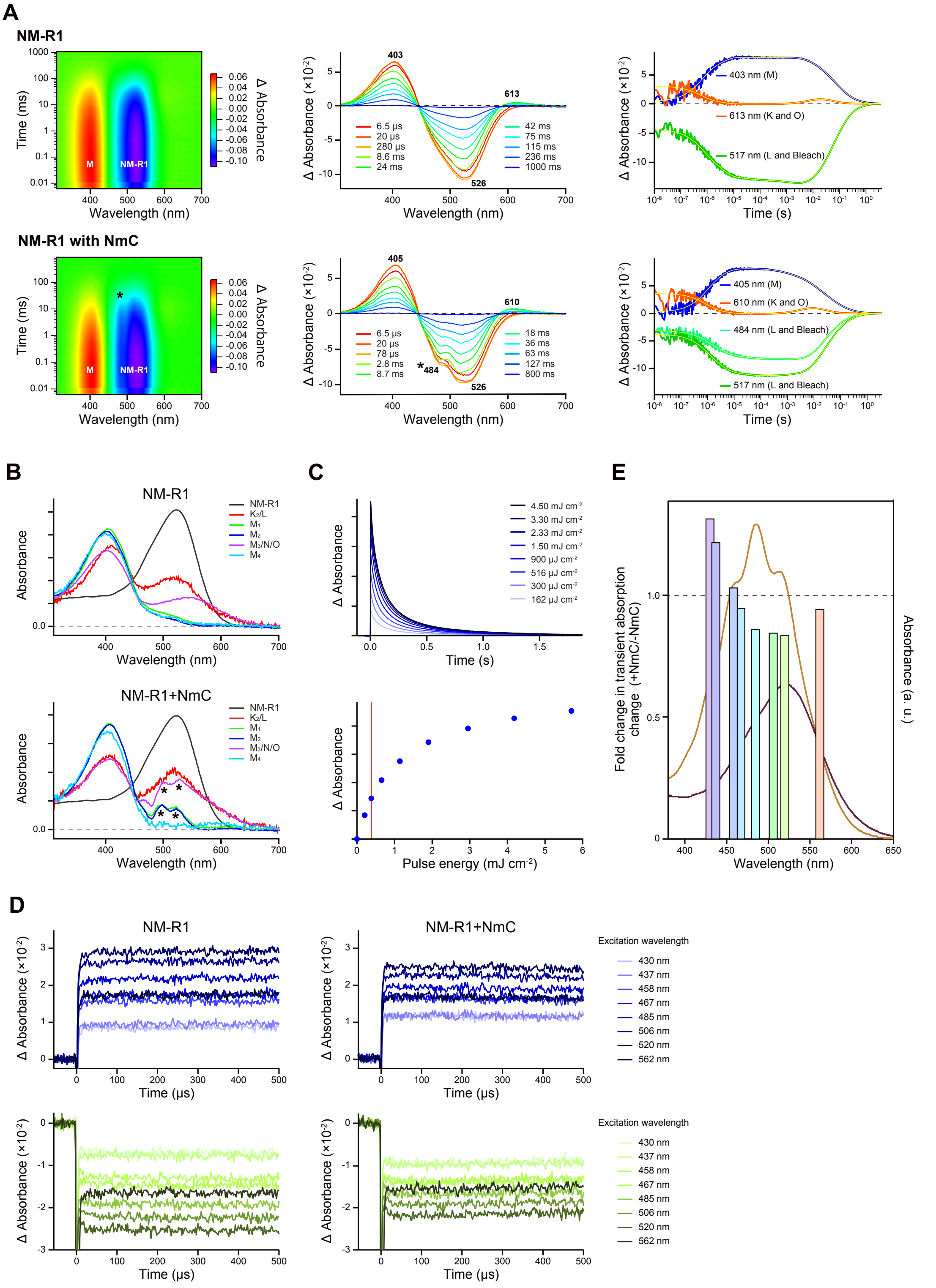


**Figure S2 | The photocycles of NM-R1 and NM-R1–NmC complex, related to Figure 3.**

(A) Two-dimensional plot of transient absorption change (left), transient absorption spectra at different time points (middle), and time courses of the transient absorption change (right) of NM-R1 without (top) and with (bottom) NmC. The multi-exponential fitting lines were indicated by yellow lines. Peaks derived from the absorption change of carotenoids are indicated by asterisks.

(B) Absorption spectra of the photointermediates of NM-R1 without (top) and with (bottom) NmC. Peaks derived from the absorption change of carotenoids are indicated by asterisks.

(C) Excitation light-energy dependence of transient absorption signal. The red line shows the excitation light intensity (0.38 mJ $cm^{-2}$) of EET measurement.

(D) Transient absorption signals probed at 390 (top) and 560 (bottom) nm representing the M accumulation and initial state bleaching, respectively, of NM-R1 (left) and NM-R1–NmC (right) excited at different wavelengths (430, 437, 458, 467, 485, 506, 520, and 562 nm).

(E) The ratios of transient absorption change in NM-R1 with and without NmC at different excitation wavelengths (430, 437, 458, 467, 485, 506, 520, and 562 nm) (bars colored according to the color of excitation light). The absorption spectra of NM-R1 without (purple line) and with (orange line) NmC were overlaid. The dashed line indicates no difference between without and with NmC.

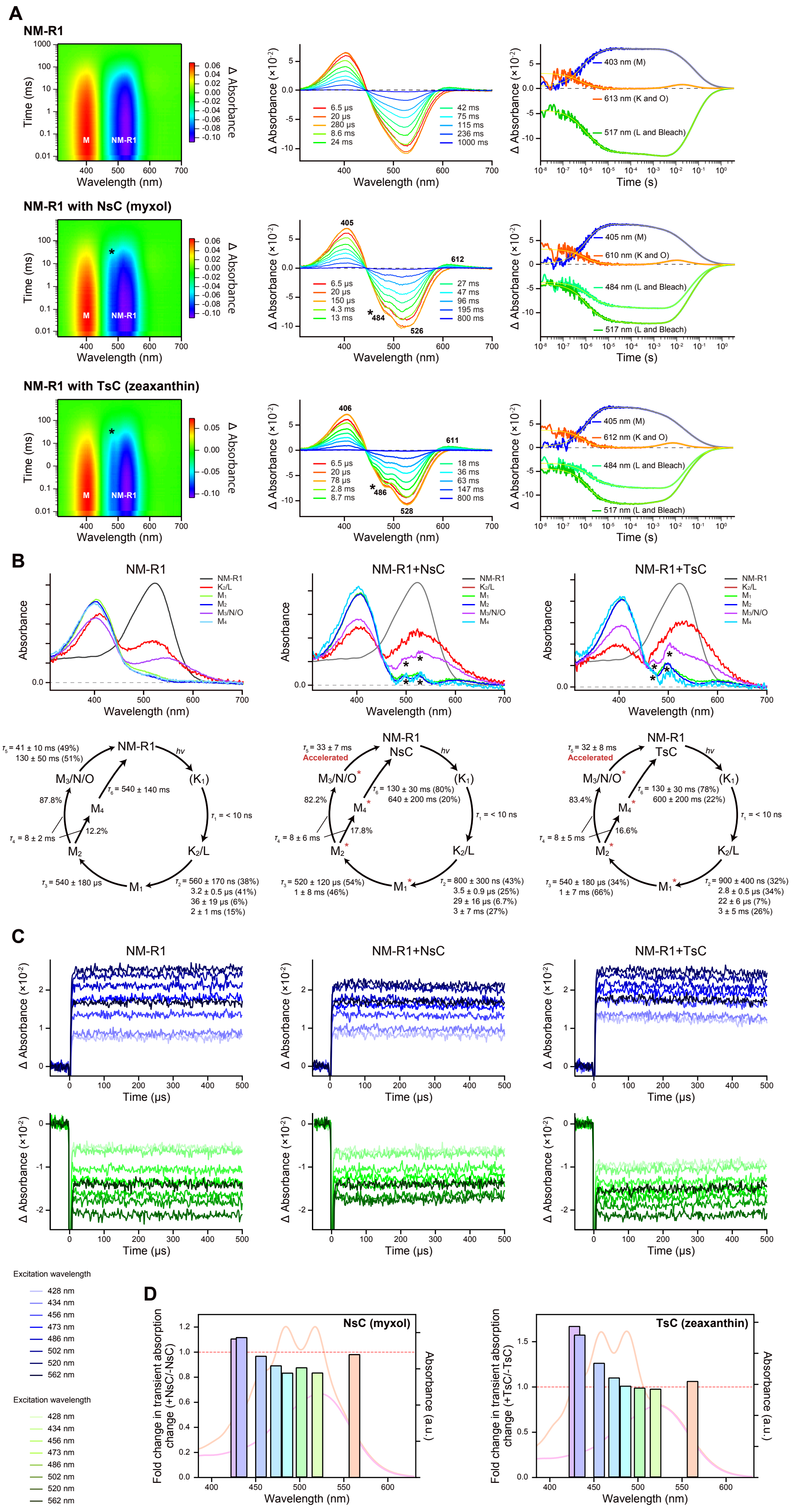
A
NM-R1
NM-R1 with NsC (myxol)
NM-R1 with TsC (zeaxanthin)
Time (ms)
Wavelength (nm)
Δ Absorbance
Δ Absorbance (×10⁻²)
Time (s)
403 nm (M)
613 nm (K and O)
517 nm (L and Bleach)
405 nm (M)
610 nm (K and O)
484 nm (L and Bleach)
517 nm (L and Bleach)
405 nm (M)
612 nm (K and O)
484 nm (L and Bleach)
517 nm (L and Bleach)
B
NM-R1
NM-R1+NsC
NM-R1+TsC
Absorbance
Wavelength (nm)
NM-R1
K2/L
M1
M2
M3/N/O
M4
C
NM-R1
NM-R1+NsC
NM-R1+TsC
Δ Absorbance (×10⁻²)
Time (μs)
Excitation wavelength
428 nm
434 nm
456 nm
473 nm
486 nm
502 nm
520 nm
562 nm
D
NsC (myxol)
TsC (zeaxanthin)
Fold change in transient absorption change (+NsC/-NsC)
Fold change in transient absorption change (+TsC/-TsC)
Absorbance (a.u.)
Wavelength (nm)

**Figure S3 |** The photocycles of NM-R1–NsC and NM-R1–TsC complex, related to Figure 3.

(A) Two-dimensional plot of transient absorption change (left), transient absorption spectra at different time points (middle), and time courses of the transient absorption change (right) of NM-R1 without carotenoids (top), with NsC (middle), and with TsC (bottom). The multi-exponential fitting lines were indicated by yellow lines. Peaks derived from the absorption change of carotenoids are indicated by asterisks.

(B) Absorption spectra of the photointermediates (top) and photocycle models (bottom) of NM-R1 without carotenoids (left), with NsC (middle), and with TsC (right). Peaks derived from the absorption change of carotenoids are indicated by asterisks.

(C) Transient absorption signals probed at 393 (top) and 560 (bottom) nm representing the M accumulation and initial state bleaching, respectively, of NM-R1 (left) and NM-R1–NsC (middle), and NM-R1–TsC (right) excited at different wavelengths (428, 434, 456, 473, 486, 502, 520, and 562 nm).

(D) The ratios of transient absorption change in NM-R1 with NsC (left) and TsC (right) at different excitation wavelengths (428, 434, 456, 473, 486, 502, 520, and 562 nm) (bars colored according to the color of excitation light). The absorption spectra of NM-R1 without (purple line) and with (orange line) carotenoids were overlaid. The dashed line indicates no difference between without and with carotenoids.

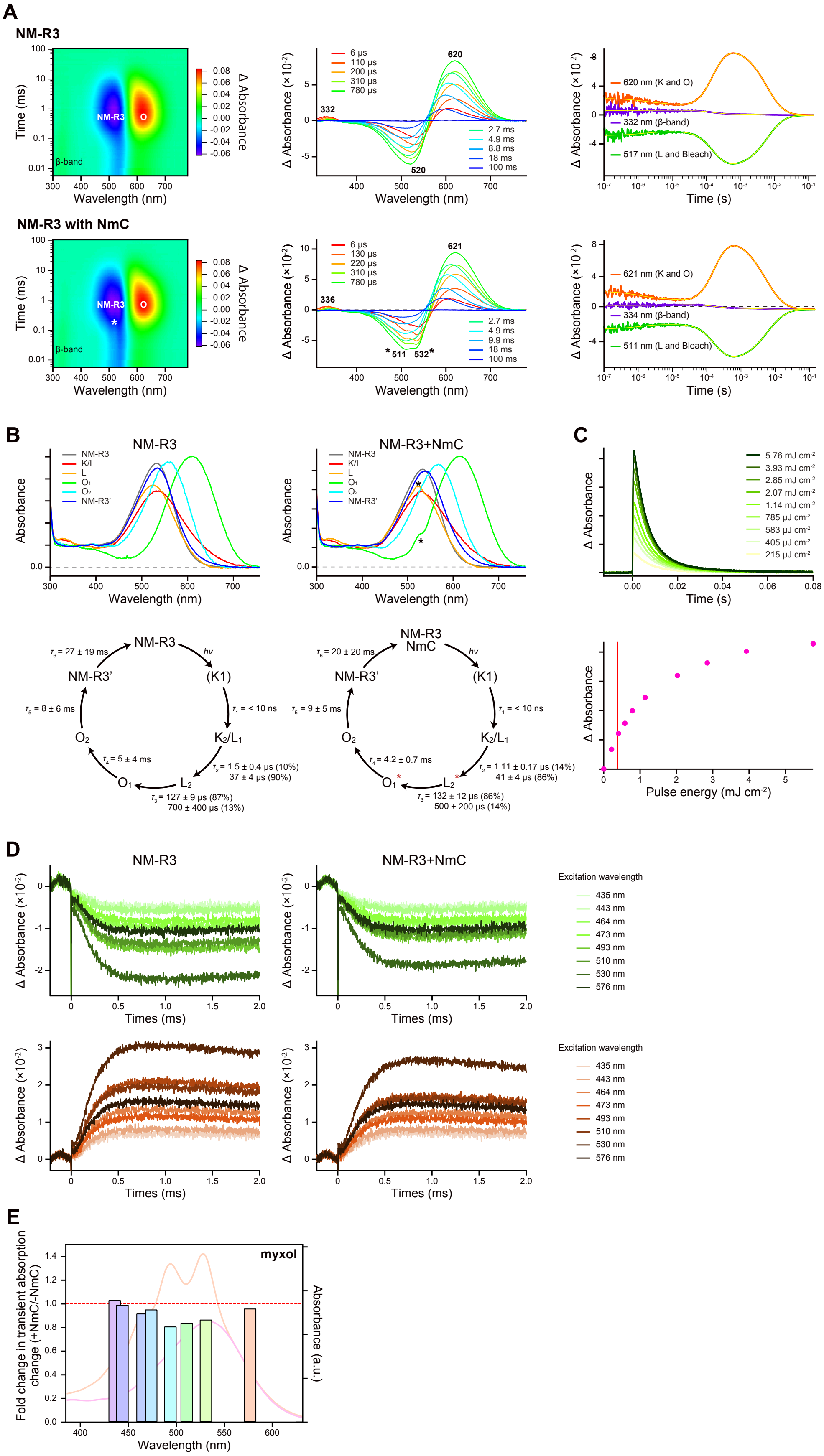
A
NM-R3
NM-R3 with NmC
Time (ms)
Wavelength (nm)
Δ Absorbance
β-band
NM-R3
O
Δ Absorbance (×10⁻²)
6 μs
110 μs
200 μs
310 μs
780 μs
2.7 ms
4.9 ms
8.8 ms
18 ms
100 ms
332
520
620
Time (s)
620 nm (K and O)
332 nm (β-band)
517 nm (L and Bleach)
130 μs
220 μs
9.9 ms
336
511
532
621
621 nm (K and O)
334 nm (β-band)
511 nm (L and Bleach)
B
NM-R3
NM-R3+NmC
Absorbance
K/L
L
O₁
O₂
NM-R3'
C
5.76 mJ cm⁻²
3.93 mJ cm⁻²
2.85 mJ cm⁻²
2.07 mJ cm⁻²
1.14 mJ cm⁻²
785 μJ cm⁻²
583 μJ cm⁻²
405 μJ cm⁻²
215 μJ cm⁻²
hν
(K1)
K₂/L₁
L₂
τ₁ = < 10 ns
τ₂ = 1.5 ± 0.4 μs (10%)
37 ± 4 μs (90%)
τ₃ = 127 ± 9 μs (87%)
700 ± 400 μs (13%)
τ₄ = 5 ± 4 ms
τ₅ = 8 ± 6 ms
τ₆ = 27 ± 19 ms
NM-R3
NmC
τ₂ = 1.11 ± 0.17 μs (14%)
41 ± 4 μs (86%)
τ₃ = 132 ± 12 μs (86%)
500 ± 200 μs (14%)
τ₄ = 4.2 ± 0.7 ms
τ₅ = 9 ± 5 ms
τ₆ = 20 ± 20 ms
Pulse energy (mJ cm⁻²)
D
Times (ms)
Excitation wavelength
435 nm
443 nm
464 nm
473 nm
493 nm
510 nm
530 nm
576 nm
E
myxol
Fold change in transient absorption change (+NmC/-NmC)
Absorbance (a.u.)

**Figure S4 |** The photocycles of NM-R3 and NM-R3–NmC complex, related to Figure 3.

(A) Two-dimensional plot of transient absorption change (left), transient absorption spectra at different time points (middle), and time course of the transient absorption change (right) of NM-R3 without (top) and with (bottom) NmC. The multi-exponential fitting lines were indicated by yellow lines. Peaks derived from the absorption change of NmC are indicated by an asterisk.

(B) Absorption spectra of the photointermediates (top) of NM-R3 without (left) and with (right) NmC. Photocycle models (bottle) of NM-R3 without (left) and with (right) NmC. The conformational change of NM-R1 affects the structure of NmC from the $L_2$ to $O_1$. Peaks derived from the absorption change of NmC are indicated by an asterisk.

(C) Dependence of transient absorption signal on excitation light intensity. The red line shows the excitation light intensity (0.38 mJ cm$^{-2}$) of EET measurement.

(D) Transient absorption signals probed at 500 (top) and 620 (bottom) nm representing initial state bleaching and the O accumulation, respectively, of NM-R3 (left) and NM-R3–NmC (right) excited at different wavelengths (435, 443, 464, 473, 493, 510, 530, and 576 nm).

(E) The ratios of transient absorption change in NM-R3 with and without NmC at different excitation wavelengths (435, 443, 464, 473, 493, 510, 530, and 576 nm) (bars colored according to the color of excitation light). The absorption spectra of NM-R3 without (purple line) and with (orange line) NmC were overlaid. The dashed line indicates no difference between without and with myxol.

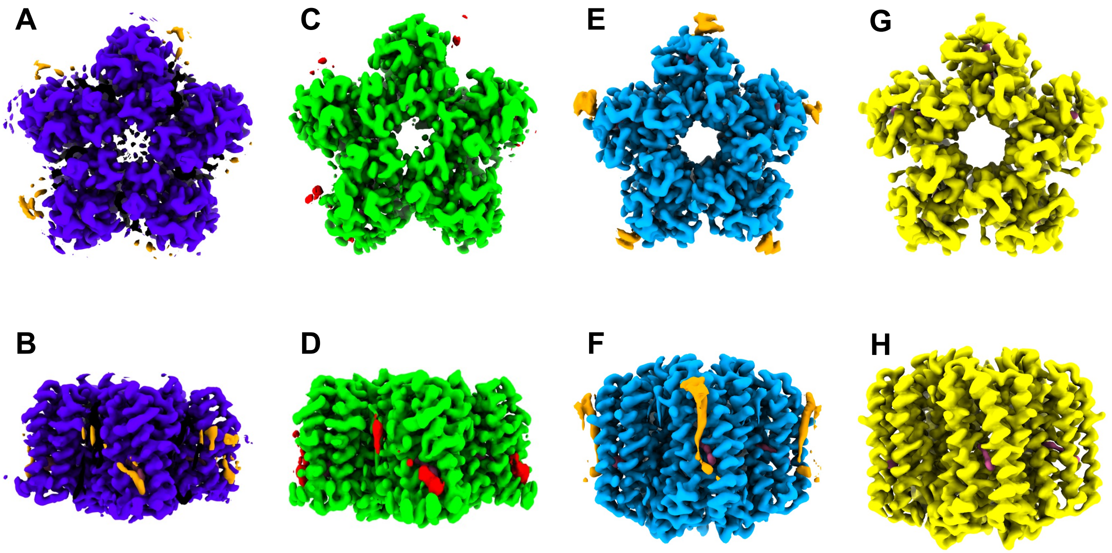


**Figure S5 |** Cryo-EM density map shown in four proteins viewed from the extracellular side of the membrane and viewed from the side, related to Figure 4 and Figure 5.

(A–B) NM-R1 bound the myxol.

(C–D) NM-R1 bound the zeaxanthin.

(E–F) NM-R3 bound the myxol.

(G–H) NM-R3.

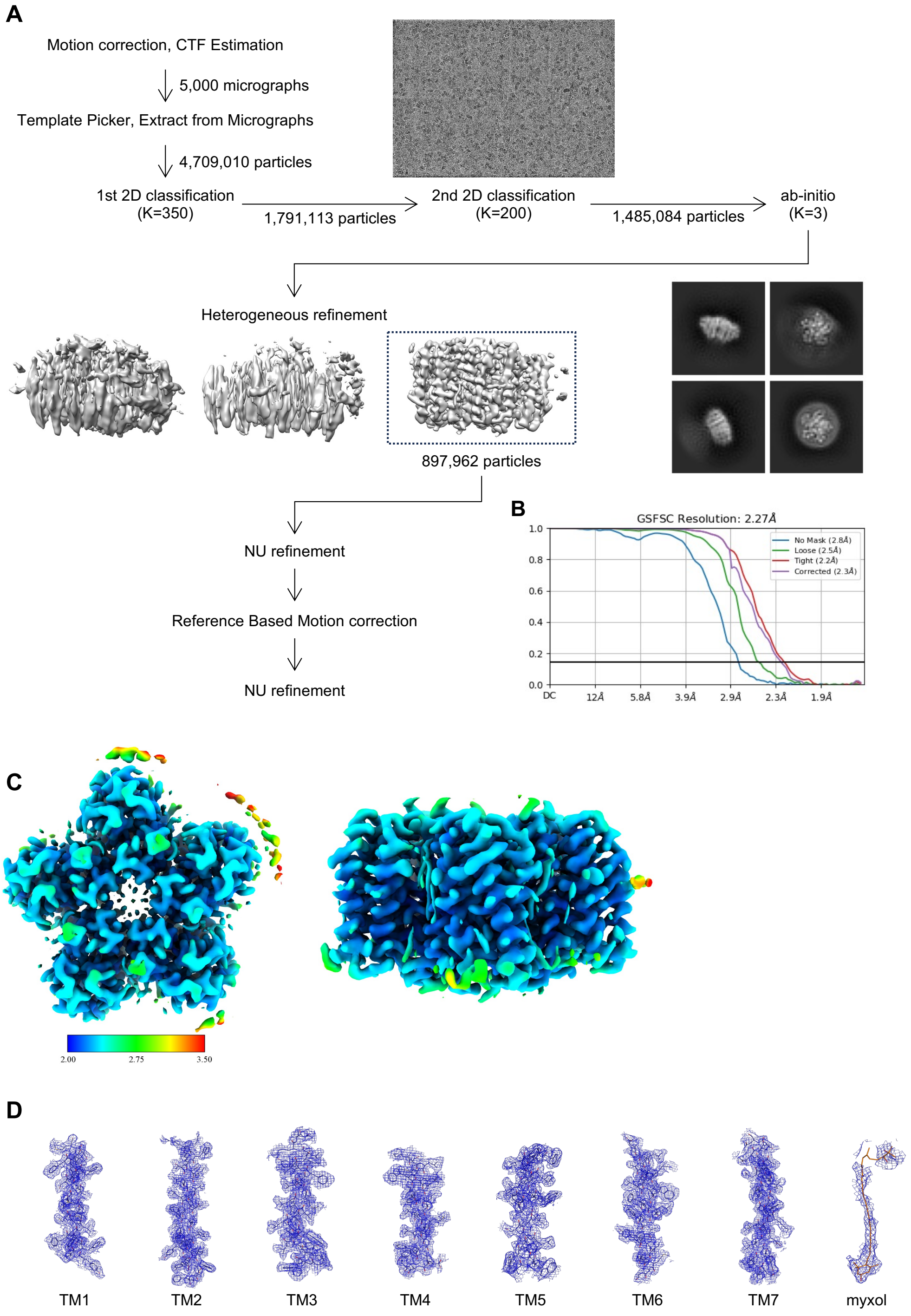


**Figure S6 |** Structural feature of NM-R1 bound the myxol, related to Figure 4.

(A) Cryo-EM single-particle analysis of the myxol-bound NM-R1.

(B) Gold-standard FSC curve used for global-resolution extimates within cryoSPARC.

(C) Local-resolution of reconstructed map as determined within cryoSPARC.

(D) Close-up view of map and side-chains density for transmembrane helices and myxol.

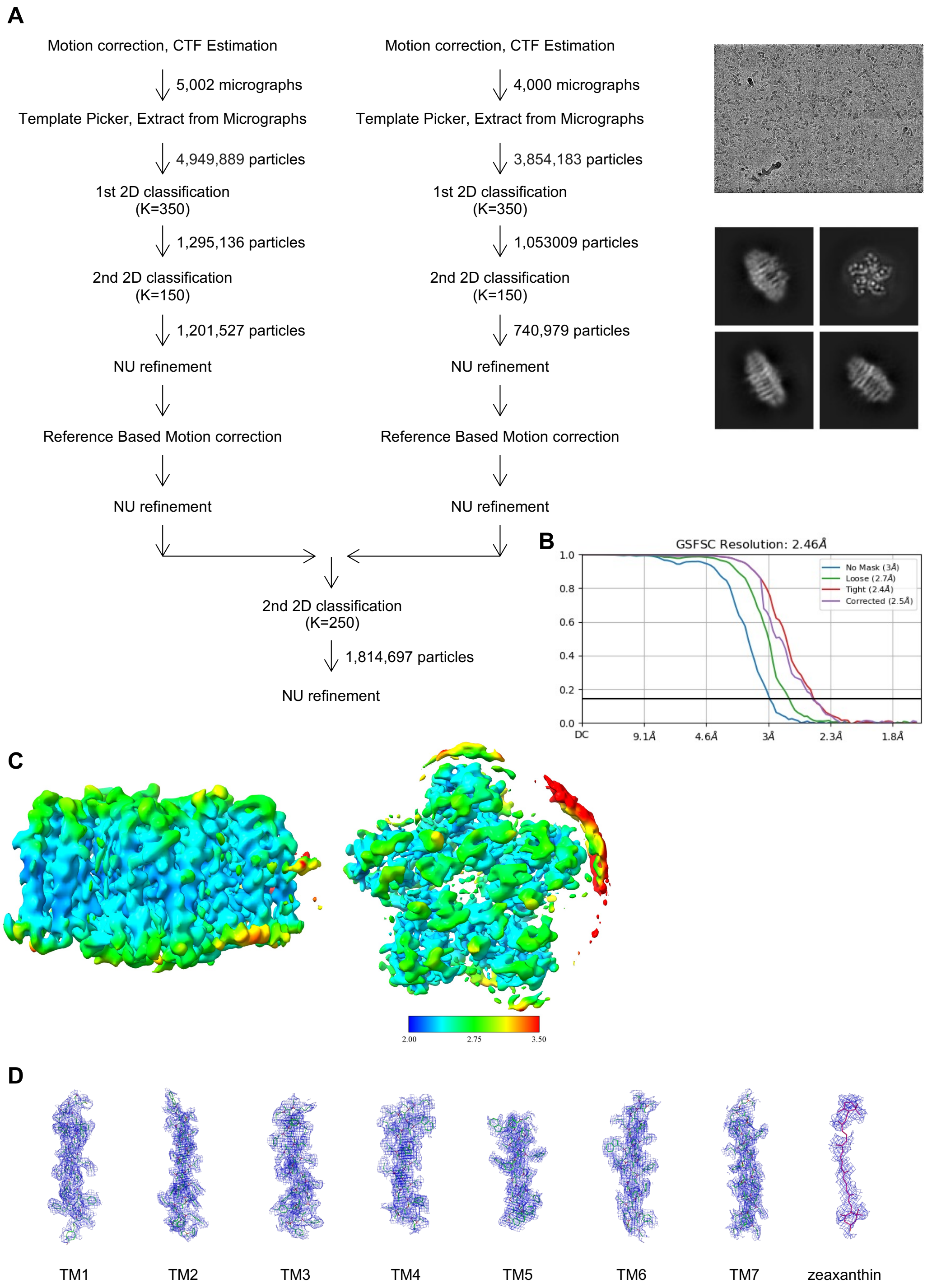


**Figure S7 |** Structural feature of NM-R1 bound the zeaxanthin, related to Figure 4.

(A) Cryo-EM single-particle analysis of the zeaxanthin-bound NM-R1.

(B) Gold-standard FSC curve used for global-resolution extimates within cryoSPARC.

(C) Local-resolution of reconstructed map as determined within cryoSPARC.

(D) Close-up view of map and side-chains density for transmembrane helices and zeaxanthin.

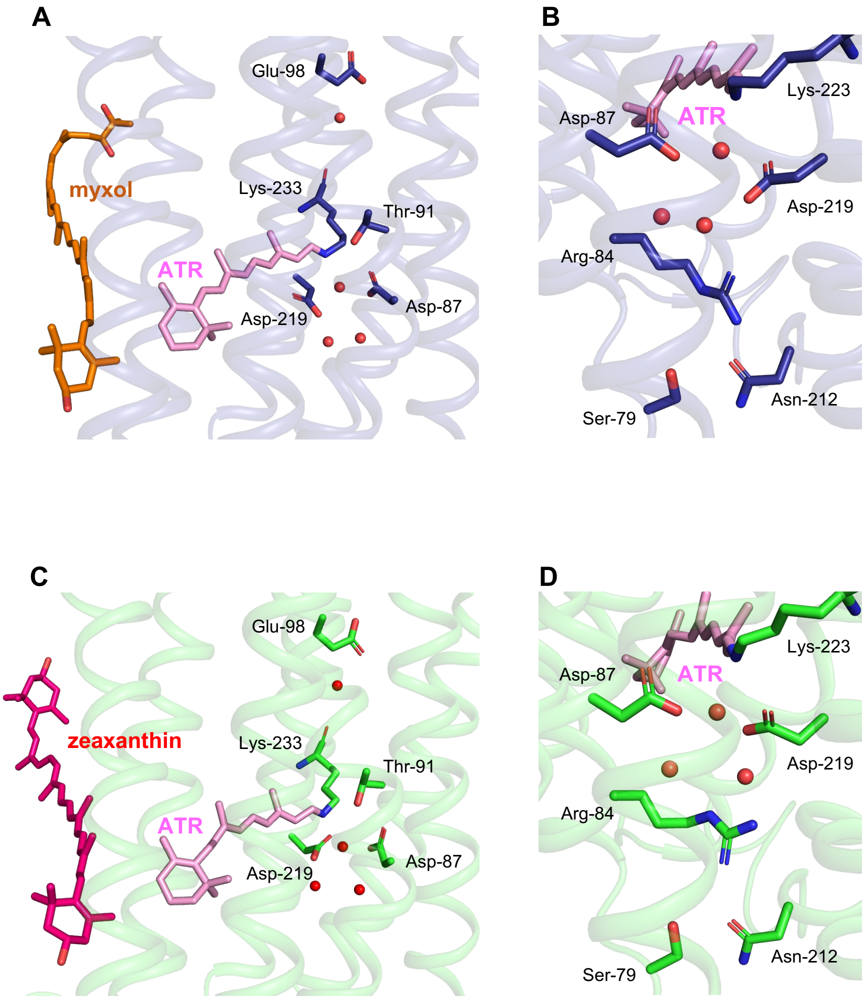


**Figure S8 |** Structure of the proton translocation pathway of NM-R1 bound the myxol or zeaxanthin, related to Figure 4.

(A) The region near the putative proton influx region, retinal, and myxol of NM-R1.

(B) Close-up views of the extracellular region in NM-R1. Stick models of key amino acids, all-*trans* retinal (ATR), and myxol are shown in blue, pink, and orange, respectively. The water ions are depicted by red sphere.

(C) The region near the putative proton influx region, retinal, and zeaxanthin of NM-R1.

(D) Close-up views of the extracellular region in NM-R1. Stick models of key amino acids, all-*trans* retinal (ATR), and zeaxanthin are shown in light green, pink, and red, respectively. The water ions are depicted by red sphere.

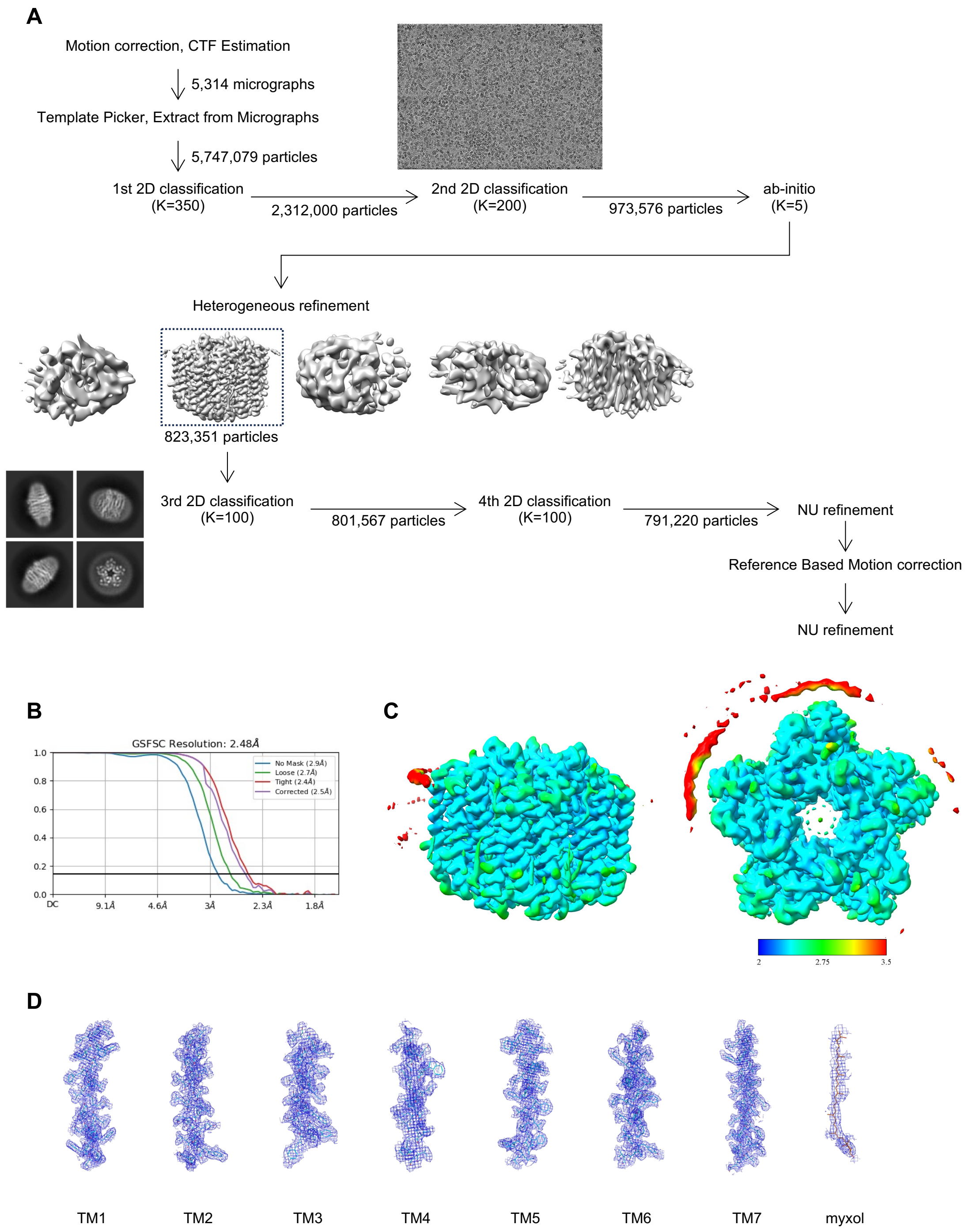


**Figure S9 |** Structural feature of NM-R3 bound the myxol, related to Figure 5.

(A) Cryo-EM single-particle analysis of the myxol-bound NM-R3.

(B) Gold-standard FSC curve used for global-resolution extimates within cryoSPARC.

(C) Local-resolution of reconstructed map as determined within cryoSPARC.

(D) Close-up view of map and side-chains density for transmembrane helices and myxol.

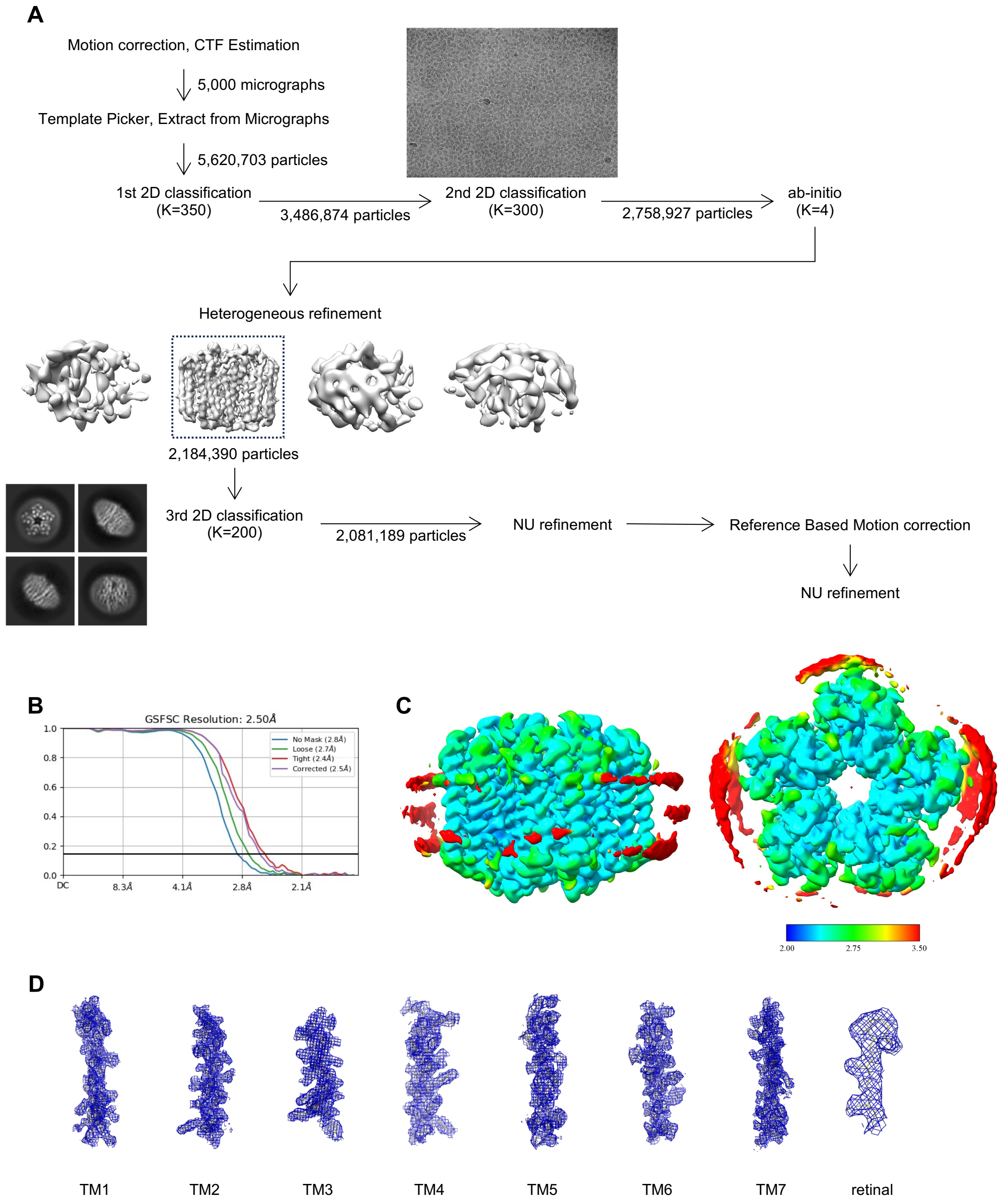


**Figure S10 |** Structural feature of NM-R3, related to Figure 5.

(A) Cryo-EM single-particle analysis of the NM-R3.

(B) Gold-standard FSC curve used for global-resolution extimates within cryoSPARC.

(C) Local-resolution of reconstructed map as determined within cryoSPARC.

(D) Close-up view of map and side-chains density for transmembrane helices and retinal.

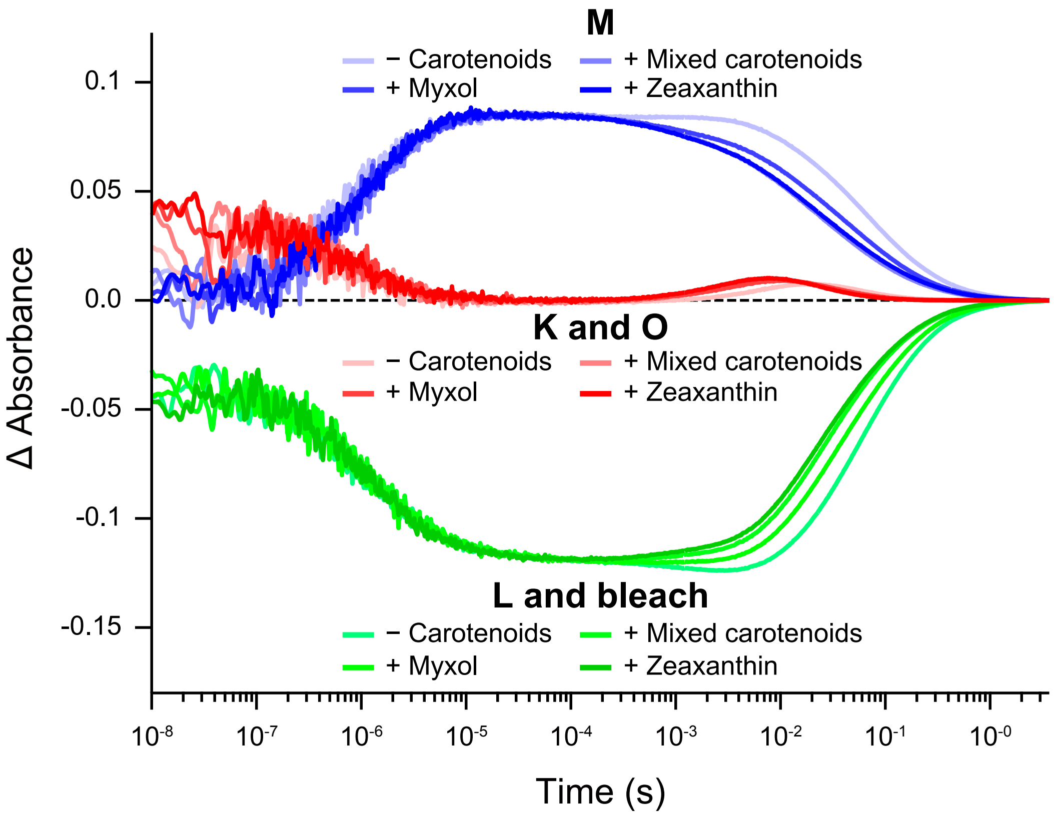


**Figure S11 | Comparison of time course of the transient absorption change of NM-R1** without carotenoids, with NmC, TsC, and NsC, related to Figure 3.

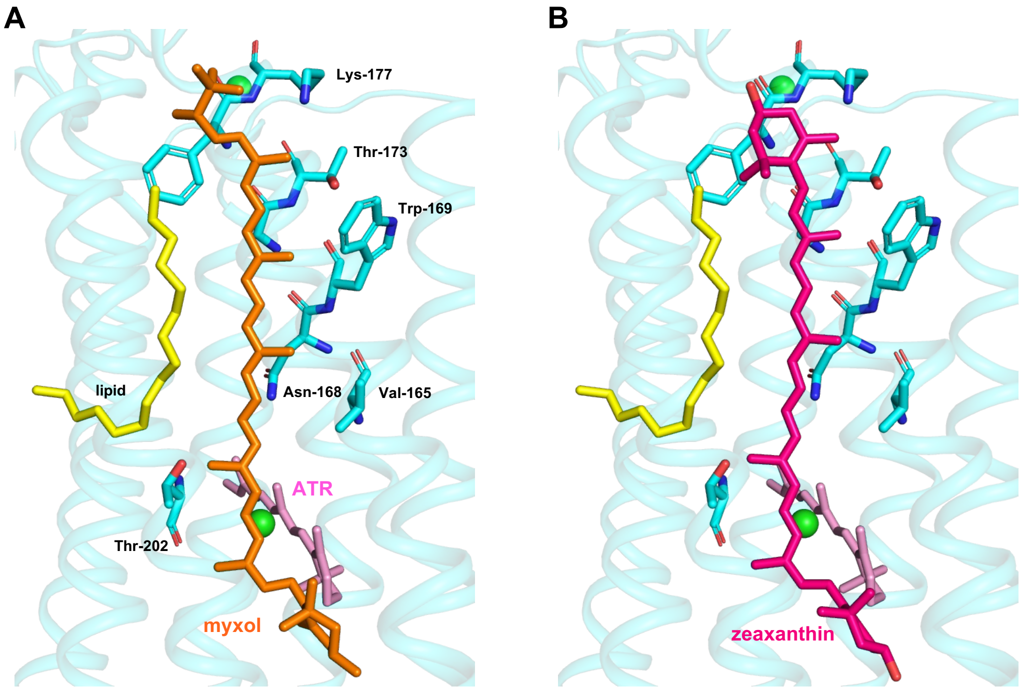


**Figure S12 |** Close up view of the myxol binding site of NM-R3, related to Figure 5.

(A) Close up view of the myxol binding site of NM-R3 between TM5 and 6.

(B) Model diagram with zeaxanthin temporarily placed where myxol exists. Stick models of retinal, myxol, zeaxanthin, and a part of acyl chain of lipid are shown in pink, orange, red, and yellow, respectively. The $Cl^{-}$ ion is depicted by light green sphere.

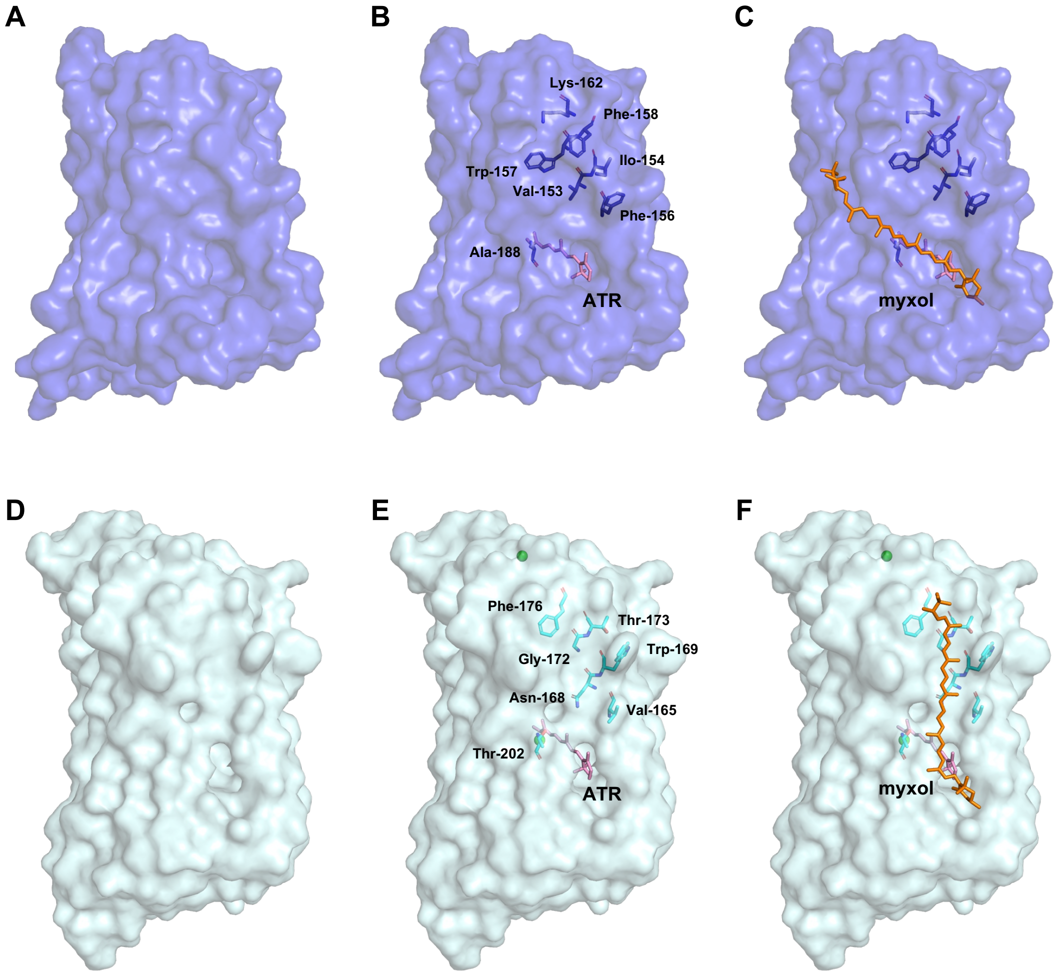


**Figure S13 |** Myxol binding site of NM-R1 or NM-R3, related to Figure 4 and Figure 5.

(A) NM-R1 surface map.

(B) NM-R1 surface map and stick model of the retinal with some of the TM5–TM6 amino acids.

(C) Further stick model of the myxol in figure (B).

(D) NM-R3 surface map.

(E) NM-R3 surface map and stick model of the retinal with some of the TM5–TM6 amino acids.

(F) Further stick model of the myxol in figure (E). Stick models of retinal and myxol are shown in pink and orange, respectively. The $Cl^-$ ion is depicted by light green sphere.

**Table. S1 |** $^{1}$H-NMR data of myxol and zeaxanthin in $CDCl_3$, related to Figure 2.

| Protons | Zeaxanthin | Myxol | Protons | Zeaxanthin | Myxol |
|---|---|---|---|---|---|
| 2α | 1.77 md | 1.77 md | 2’ | | 4.00 dd |
| 2β | 1.48 dd | 1.48 dd | 2’-OH | | 2.08 d |
| 3 | 4.00 m | 4.00 dd | 3’ | | 5.71 dd |
| 3-OH | 1.37 d | 1.37 d | 4’ | | 6.41 d |
| 4α | 2.39 dd | 2.39 dd | 6’ | | 6.21 d |
| 4β | 2.05 dd | 2.05 dd | 7’ | | 6.58 d |
| 7 | 6.11 | 6.11 | 8’ | | 6.39 d |
| 8 | 6.16 d | 6.16 d | 10’ | | 6.19 d |
| 10 | 6.16 d | 6.16 d | 11’ | | 6.63 m |
| 11 | 6.63 m | 6.63 m | 12’ | | 6.39 d |
| 12 | 6.36 d | 6.36 d | 14’ | | 6.28 d |
| 14 | 6.26 d | 6.26 d | 15’ | | 6.63 m |
| 15 | 6.63 m | 6.63 m | 16’ | | 1.242 s |
| 16 | 1.073 s | 1.074 s | 17’ | | 1.181 s |
| 17 | 1.073 s | 1.074 s | 18’ | | 1.927 s |
| 18 | 1.738 s | 1.738 s | 19’ | | 1.979 s |
| 19 | 1.969 s | 1.972 s | 20’ | | 1.987 s |
| 20 | 1.974 s | 1.972 s | | | |

Chemical shifts, δ (ppm) and multiplicity (d, doublet; m, multiplet; s, singlet).

**Table. S2 |** Cryo-EM data collection, refinement, and validation statistics, related to Figure 4 and Figure 5.

| | NM-R1-myxol | NM-R1-zeaxanthin | NM-R3-myxol | NM-R3 |
|---|---|---|---|---|
| PDB ID | 9JOV | 9JOU | 9JOW | 9JOX |
| EMDB ID | EMD-61686 | EMD- 61685 | EMD-61687 | EMD-61688 |
| **Data collection and processing** | | | | |
| Magnification | 105,000 | 105,000 | 105,000 | 105,000 |
| Voltage (kV) | 300 | 300 | 300 | 300 |
| Electron exposure ($e^-$/Å$^2$) | 60.425 | 54.0 | 60.425 | 57.534 |
| Defocus range (μm) | −0.8 to −2.0 | −0.8 to −2.0 | −0.8 to −2.0 | −0.8 to −2.0 |
| Pixel size (Å) | 0.83 | 0.83 | 0.83 | 0.83 |
| Symmetry | C1 | C1 | C1 | C1 |
| No. of movie | 5,000 | 9,002 | 5,314 | 5,000 |
| No. of initial particle images | 4,709,010 | 8,804,012 | 5,747,079 | 5,620,703 |
| No. of final particle images | 897,953 | 1,814,697 | 791,219 | 2,081,189 |
| Map resolution (Å) | 2.27 | 2.46 | 2.48 | 2.50 |
| FSC threshold | 0.143 | 0.143 | 0.143 | 0.143 |

| **Refinement** | | | | |
|---|---|---|---|---|
| Model resolution (Å) | 2.27 | 2.46 | 2.48 | 2.50 |
| Model composition | | | | |
| No. of non-hydrogen atoms | 10,041 | 9,913 | 11,677 | 11,249 |
| No. of protein residues | 1,150 | 1,150 | 1,320 | 1,320 |
| B factors ($Å^2$) | | | | |
| Protein | 85.91 | 120.57 | 94.61 | 113.47 |
| R.m.s. deviations | | | | |
| Bond lengths (Å) | 0.011 | 0.010 | 0.014 | 0.013 |
| Bond angles (°) | 1.794 | 1.493 | 2.083 | 1.976 |
| Validation | | | | |
| MolProbity score | 1.99 | 2.30 | 1.74 | 2.16 |
| Clash score | 9.17 | 14.93 | 5.20 | 5.90 |
| Poor rotamers (%) | 4.59 | 6.56 | 3.52 | 5.19 |
| Ramachandran plot | | | | |
| Favored (%) | 98.51 | 98.68 | 97.79 | 95.73 |
| Allowed (%) | 1.49 | 1.32 | 2.21 | 4.27 |
| Disallowed (%) | 0 | 0 | 0 | 0 |
